# Deconvolution of substrate processing by the 26S proteasome reveals a selective kinetic gateway to degradation

**DOI:** 10.1101/359695

**Authors:** J.A.M. Bard, C. Bashore, K.C. Dong, A. Martin

## Abstract

The 26S proteasome is the principle macromolecular machine responsible for protein degradation in eukaryotes. However, little is known about the detailed kinetics and coordination of the underlying substrate-processing steps of the proteasome, and their correlation with observed conformational states. Here, we used reconstituted 26S proteasomes with unnatural amino acid-attached fluorophores in a series of FRET and anisotropy-based assays to probe substrate-proteasome interactions, the individual steps of the processing pathway, and the conformational state of the proteasome itself. We develop a complete kinetic picture of proteasomal degradation, which reveals that the engagement steps prior to substrate commitment are fast relative to subsequent deubiquitination, translocation and unfolding. Furthermore, we find that non-ideal substrates are rapidly rejected by the proteasome, which thus employs a kinetic proofreading mechanism to ensure degradation fidelity and substrate prioritization.

## Main Text

Specific protein degradation is essential for quality control, homeostasis, and the regulation of diverse cellular processes, such as the cell cycle and stress response*(1, 2)*. In eukaryotic cells, this degradation is primarily catalyzed by the 26S proteasome, which recognizes, unfolds, and degrades target proteins that have been modified with lysine-attached poly-ubiquitin chains*(3)*. To accomplish its dual roles in quality control and signaling, the proteasome must be able to degrade certain proteins consistently and under tight kinetic constraints, while maintaining a high promiscuity necessary to process thousands of polypeptides with diverse characteristics, yet avoiding unregulated proteolysis of cellular proteins in general. This precise regulation is accomplished by the proteasome’s complex molecular architecture and a bipartite degradation signal for substrate recognition that consists of a suitable poly-ubiquitin chain and an unstructured initiation region of sufficient length and appropriate sequence composition*(4-9)*. The proteolytic active sites of the proteasome are sequestered inside a barrel shaped compartment termed the 20S core particle, which excludes folded proteins and even large unfolded polypeptides*(10)*. Substrate access to these active sites is controlled by the 19S regulatory particle that caps one or both ends of the 20S core, recruits ubiquitinated proteins, unfolds them, and translocates the unstructured polypeptides through a central pore into the 20S core for proteolytic cleavage. This regulatory particle can be further divided into two 9-subunit subcomplexes, the base and lid*(11)*. The base contains multiple ubiquitin receptors and a ring-shaped AAA+ (ATPases Associated with diverse cellular Activities) motor formed by the six distinct ATPases Rpt1-Rpt6*(12-16)*. Each ATPase subunit consists of an N-terminal helix, an OB-fold domain, and a C-terminal AAA+ motor domain. In the heterohexamer, the N-terminal helices of neighboring Rpt-subunit pairs engage in coiled-coil interactions, while the six OB-fold domains form a rigid N-ring that sits atop the ATPase ring. After ubiquitin binding, a substrate’s flexible initiation region likely must reach through the N-ring before engaging with conserved pore loops of the ATPase domains for mechanical pulling that is driven by ATP binding and hydrolysis*(17, 18)*. The lid subcomplex is bound to one side of the base and contains the deubiquitinating enzyme Rpn11. In the proteasome holoenzyme, Rpn11 is positioned near the entrance to the AAA+ motor, allowing it to remove ubiquitin chains *en bloc* from substrates during translocation*(19-23)*.

Both *in vivo* and *in vitro* structural studies revealed that the 26S proteasome adopts multiple conformations, which can be divided into a substrate-free apo (s1) state and substrate-processing (s3-like) states*(24-31)*. In the s1 state, the primary conformation adopted by the ATP-bound proteasome in the absence of protein substrate, the ATPase domains form a steep spiral staircase, and Rpn11 is offset from the central pore of the motor, which itself is not aligned with the entrance to the 20S core. The s3-like state, which can be stabilized by stalling a substrate during translocation, appears more conducive to processive degradation, as the central channel of the motor aligns with the N-ring and the entrance to the 20S core, and it contains a more planar ATPase ring in which neighboring Rpt subunits form uniform interfaces*(30)*. In addition, during the transition from the s1 to s3 states, the lid rotates relative to the base, placing Rpn11 in a position directly above the pore entrance, where it is able to remove ubiquitin from a substrate polypeptide as it translocates into the motor. Similar s3-like states have also been observed by incubating the proteasome with nucleotide analogs such as ATPγS*(27)*.

Based on the available structural and biochemical information, we can infer the processing steps necessary for substrate degradation, which include the binding of ubiquitin to a proteasomal receptor, insertion of the unstructured initiation region into the pore of the ATPase motor, the removal of ubiquitin chains by Rpn11, and the unfolding and translocation of the polypeptide into the 20S core for proteolysis. However, very little is known about the relative timing and coordination of these events, and the rate-limiting step of proteasomal degradation is still unclear. It also remains elusive what induces the proteasome to switch from the s1 to the s3-like states and how these distinct conformations are coordinated with the processing pathway. Here we reveal the complete kinetic picture of substrate processing by the 26S proteasome. We designed a series of fluorescence- and FRET-based assays to specifically measure the kinetics of individual degradation steps in a fully reconstituted and thus well-controlled system, without contamination by accessory proteins, like the non-essential deubiquitinase Ubp6*(32)*. These novel tools allowed us to track the interactions between a substrate’s initiation region and the AAA+ motor, the conformational changes of the proteasome, substrate deubiquitination by Rpn11, and cleavage into peptides. Our results reveal the rate-limiting step of degradation, identify the trigger for the conformational switch from the s1 to the s3 state of the proteasome, and elucidate the regulation of Rpn11’s deubiquitination activity. Furthermore, we analyzed the degradation effects of varying substrate characteristics, including the stability of the folded domain, the number of ubiquitin chains, and the length and composition of the unstructured initiation region, offering new insights into how the proteasome selects and prioritizes its substrates in a complex cellular environment.

## Unnatural amino-acid labeling of the proteasome

To deconvolute individual steps of substrate degradation and track the coupled conformational changes of the proteasome, we sought to develop Förster resonance energy transfer (FRET)-based assays sensitive to specific processing events. One requirement for these assays was the ability to site-specifically attach fluorophores to the 19S regulatory particle with minimal perturbation of its structure and function. Building on our previously established recombinant expression systems for base and lid sub-complexes of the *Saccharomyces cerevisiae* 26S proteasome and their purification from *Escherichia coli*(*18, 31*), we devised a method for the site-specific incorporation and labeling of 4-azido-L-phenylalanine (AzF)*(33)*. We introduced a highly evolved AzF synthetase on an optimized plasmid into our base and lid expression systems, allowing for the purification of unnatural amino acid-containing sub-complexes and the covalent attachment of a fluorophore at specific sites within the 19S regulatory particle (Fig. 1A)(*34, 35*). To increase the labeling specificity, solvent-exposed cysteines of the lid and base were reversibly protected before reacting the incorporated AzF with a dibenzocyclooctyne-linked fluorophore*(36)*. This procedure led to 50-80% labeling efficiency and minimal off-target reactions (Fig. 1A). Importantly, proteasomes reconstituted with these fluorescently labeled base and lid sub-complexes exhibited full activity in substrate degradation (Fig. 1B, Fig. S1).

**Fig. 1.**
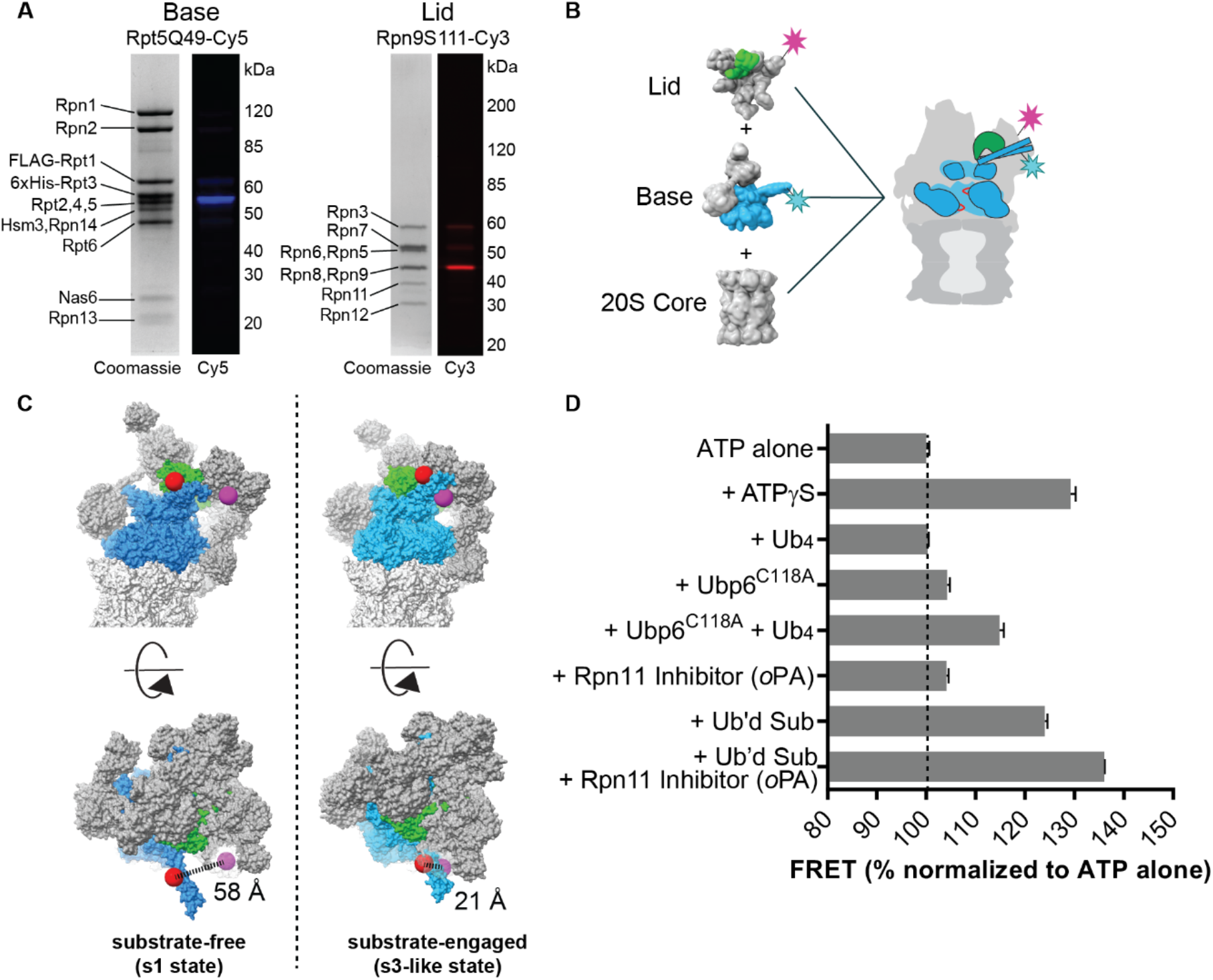
Site-specific labeling of the 26S proteasome and a steady-state assay for its conformations. **(A)** SDS-PAGE analysis of base and lid subcomplexes with AzF incorporated into Rpn5 and Rpn9, and then labeled with Cy5 and Cy3, respectively. Images of both, the Coomassie-stained gels and fluorescence detection are shown. (**B**) Schematic for the *in vitro* reconstitution of the 26S proteasome from lid subcomplex with Cy3-labeled-Rpn9, base with Cy5-labeled-Rpt5, and 20S core. (**C**) A comparison of the substrate-free and substrate-engaged states, showing that the distance between Rpt5-Q49 and Rpn9-Sl 11 changes by 37 Å during the conformational transition. The 26S proteasome is depicted with the AAA+ ATPases in blue, Rpn11 in green, and the rest of the regulatory particle as well as the 20S core in grey. Models were generated from surface representations of atomic models (PDB: 5mp9, 5mpd, 5mpb). The red and purple spheres indicate the positions of the labeled residues on Rpt5 and Rpn9, respectively. **(D)** Steady-state FRET signals between Cy3-labeled lid (RpnS111AzF-Cy3) and Cy5-labeled base (Rpt5Q49AzF-Cy5) under various conditions, normalized to the signal for proteasome with ATP alone. Ubp6^C118A^ is catalytically inactive, Ub_4_ represents a linearly fused tetra-ubiquitin, and the ubiquitinated substrate is titin-I27^V15P^ with a 35-residue tail (titin-I27^V15P^-35). Shown are the mean and s.d. for *N=3.*

## Tracking the Conformational State of the Proteasome

Published cryo-EM structures of the 26S proteasome revealed major conformational changes between the substrate-free and substrate-processing states, yet it remains unclear how these transitions are coupled to specific steps of substrate degradation. We identified the lid subunit Rpn9 and the N-terminal coiled-coil of the base subunits Rpt4 and Rpt5 as promising positions for the placement of a donor-acceptor pair to directly monitor the proteasome conformational dynamics through FRET (Fig. 1C). As the lid rotates relative to the base during the transition to a substrate-engaged state, the distance between Rpn9 and the N-terminal helix of Rpt5 decreases by almost 40 Å, and we predicted that the FRET efficiency for fluorophores at these positions would increase accordingly.

Proteasomes were thus reconstituted using lid with donor-labeled Rpn9-S111AzF and base with acceptor-labeled Rpt5-Q49AzF, and bulk FRET efficiencies were measured under steady-state conditions (Fig. 1D). Incubation with the non-hydrolyzable nucleotide analog ATPγS has previously been shown to induce the substrate engaged-like s3 state*(27)*, and it correspondingly caused a significant increase in FRET compared to proteasome in ATP alone (Fig. 1D). Interaction with ubiquitin-bound Ubp6 has also been found to shift the conformational equilibrium towards the substrate-engaged state*(37, 38)*, which was confirmed by our assay showing an increase in FRET efficiency when catalytically inactive Ubp6 was added together with tetra-ubiquitin. There was no change in FRET, however, when 100 μM unanchored ubiquitin chains were added, indicating that ubiquitin binding to proteasomal receptors does not induce a conformational change.

We then measured the FRET signal of actively degrading proteasomes and observed an increase in FRET similar to that of proteasomes trapped in the s3 state by ATPγS. The signal increased further upon addition of ubiquitinated substrate in the presence of 1,10-phenanthroline (oPA), an inhibitor of the deubiquitinating enzyme Rpn11*(19)*. Preventing Rpn11-mediated deubiquitination inhibits degradation by stalling substrate translocation through the central channel, most likely when the attached ubiquitin chain reaches the narrow entrance to the N-ring pore, thus trapping and synchronizing the proteasomes in a substrate engaged state (Fig. S1)*(21)*. We conclude that actively processing proteasomes spend most of the time in an s3-like conformation.

## Rapid substrate engagement induces the proteasome conformational switch

While the steady-state FRET-based assay confirmed that actively degrading proteasomes switch away from the s1 state, it provided no information about when this switch occurs. To determine the relationship between the conformational change and specific processing steps, and to gain insight into the overall coordination of substrate degradation, we developed a series of fluorescence-based assays for measuring detailed kinetics (Fig. 2). The model substrate for these assays consisted of a titin-I27 domain that was destabilized by a V15P mutation and fused with a C-terminal unstructured initiation region (a 35 amino acid ‘tail’, see Table S2)(*37, 39*). This titin-I27^V15P^-35 substrate also contained a single lysine to enable the attachment of a poly-ubiquitin chain in a defined position.

**Fig. 2.**
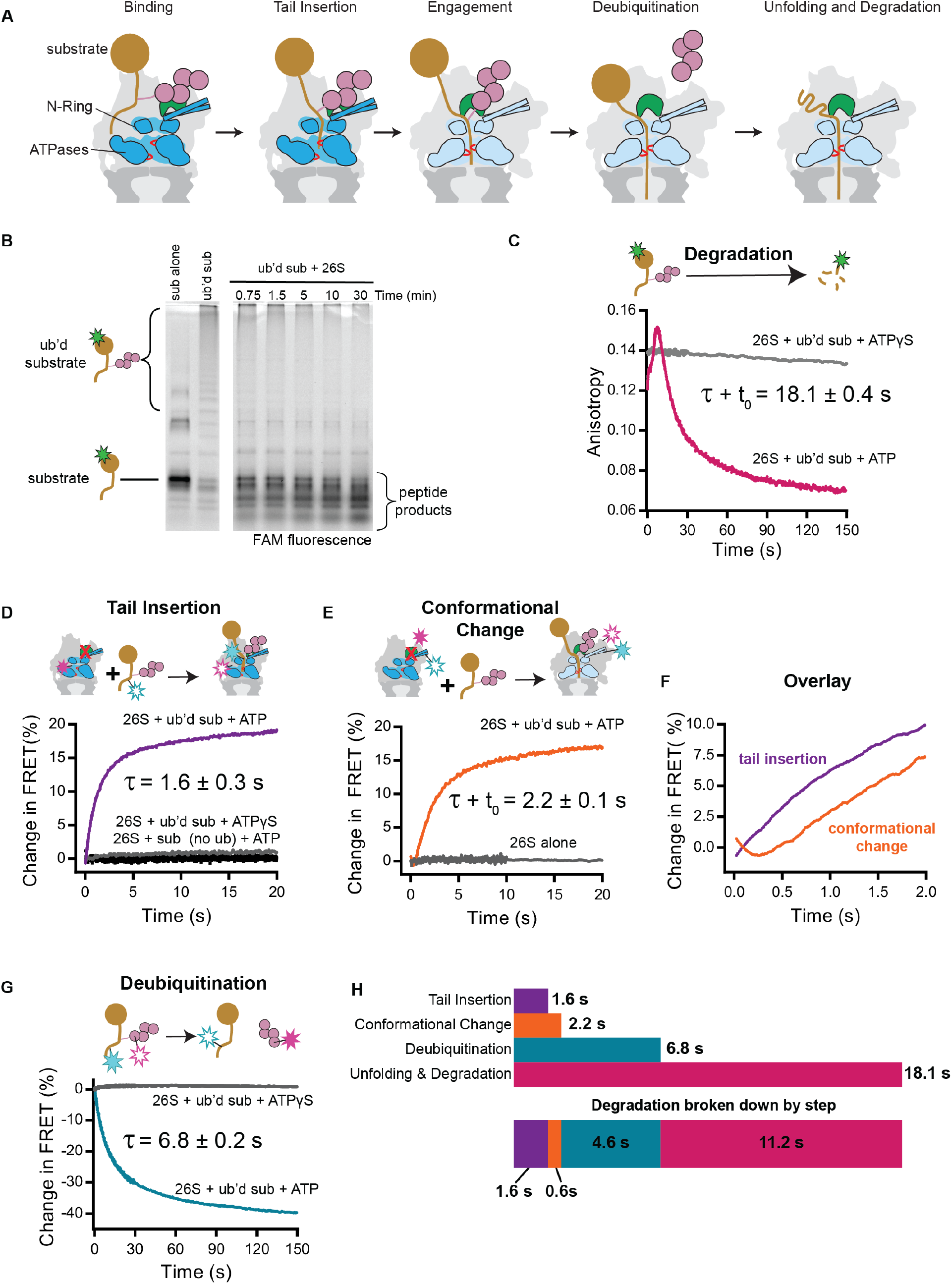
Substrate engagement triggers the conformational switch of the proteasome. **(A)** Schematic model for the substrate processing pathway. Ubiquitin chains (pink) target a substrate (gold) to bind to the 26S proteasome, which resides in the substrate-free, s1 state. The unstructured tail of a substrate is inserted into the pore of the AAA+ motor (blue), where it interacts with pore loops (red) that drive translocation. After this substrate engagement, a major conformational change rearranges the AAA+ motor and shifts Rpn11 (green) to a central position above the entrance to the pore, allowing translocation-coupled deubiquitination. Ubiquitin-chain removal is followed by mechanical substrate unfolding and threading of the polypeptide into the 20S core for proteolytic cleavage. **(B)** Single-turnover degradation of ubiquitinated 5-FAM-titin-I27^V15P^-35 by reconstituted 26S proteasome is tracked by SDS-PAGE and visualized by the fluorescence of the N-terminally attached 5-carboxyfluorescein (5-FAM, green star). **(C)** Single-turnover degradation of ubiquitinated 5-FAM-titin-I27^V15P^-35 is tracked by fluorescence anisotropy in the presence of ATP or ATPγS. The total time for degradation is derived from the sum of the time constant τ for the exponential decay of anisotropy and the time t_0_ for the initial anisotropy increase. **(D)** Substrate-tail insertion is tracked by FRET between the 26S proteasome reconstituted with Cy3-labeled base (Rpt1-I191AzF-Cy3) and the ubiquitinated titin-I27^V15P^-35 substrate modified with Cy5 on its unstructured tail. At t=0, an excess of substrate was added to proteasomes with *o*PA-inhibited Rpn11 in the presence of either ATP (purple) or ATPγS (grey). A control with non-ubiquitinated, Cy3-labeled titin-I27^V15P^-35 substrate is depicted in black. Shown is the signal of the Cy3 channel, normalized to initial fluorescence. **(E)** Conformational state of the proteasome over time, tracked by FRET between Cy5-labeled base (Rpt5Q49AzF-Cy5) and Cy3-labled lid (Rpn9S111AzF-Cy3). At t=0, an excess of ubiquitinated titin-I27^V15P^-35 or buffer was added to double-labeled proteasomes with oPA-inhibited Rpn11. Shown is the Cy5 channel, normalized to initial fluorescence. The total time for the conformational change is derived from the sum of the time constant τ for the exponential increase of FRET and the time t_0_ for the initial delay. **(F)** Overlay of fluorescence traces for tail insertion and conformational change from **D** and **E** reveal a delay of 400 ms. **(G)** Deubiquitination tracked by FRET between Cy3-labeled ubiquitin and a Cy5 label attached adjacent to the single ubiquitinated lysine in titin-I27^V15P^-35. The substrate was ubiquitinated with Cy3-labeled ubiquitin, and at t=0 mixed with excess proteasome in the presence of ATP or ATPγS. Shown is the Cy5 channel, normalized to initial fluorescence. **(H)** Time constants of the substrate-processing steps as calculated from each assay individually or after accounting for the preceding steps. All curves in **C** - **G** are representative traces, and time constants are derived from averaging fits of independent experiments, shown with s.d. (N ≥ 3).

The total time required for degradation of this substrate was determined by tracking the anisotropy of a fluorophore attached to the N-terminus of the titin folded domain(*40*). Gel-based assays confirmed the rapid degradation into peptides (Fig. 2B), and Michaelis-Menten analyses established the substrate affinity (K_M_ = 0.22 μΜ) and degradation rate (k_cat_ = 2.34 sub enz^−1^ min^−1^) under multiple-turnover, steady-state conditions (Fig. S2). In subsequent single-turnover degradation experiments with saturating concentrations of proteasome, we observed an initial fast increase in anisotropy, followed by a second increase, and an exponential signal decay (Fig. 2C, S3, Table S3). When the substrate was mixed with proteasomes containing the catalytically dead Rpn11^AXA^ mutant, the signal also rapidly increased, but did neither show the second increase nor the exponential decay of anisotropy (Fig. S3). Mixing of the substrate with ATPγS-bound wild-type proteasomes led to no change in signal (Fig. 2C). The initial anisotropy increase is thus likely a result of substrate engagement by the AAA+ motor, with a further increase seen after the deubiquitination event, as the folded domain is pulled towards the pore. The decay then reflects the mechanical unfolding, translocation, and cleavage of the substrate into small peptides. This decay (and all kinetic traces later measured for the individual substrate-processing steps) fit best to a double exponential curve, consisting of a dominant fast phase and a low-amplitude slow phase. Since the slow-phase likely originates from a small population of partially aggregated or incompletely ubiquitinated substrate, we focused our analyses on the fast phase, as done in previous studies(*18, 21*). Combining the time constant for the exponential decay in anisotropy (τ = 11 s) and the delay associated with the initial increase (t_0_ = 7 s) reveals that complete degradation of our titin model substrate occurs with a time constant of 18 s. The first step of substrate processing after ubiquitin binding requires the insertion of the unstructured initiation region into the pore of the proteasomal motor(*9*). We specifically tracked the kinetics of this tail-insertion process using a FRET-based assay that relied on the energy transfer between a donor fluorophore placed near the central channel of the motor, in the linker between the N-domain and the ATPase domain of Rpt1 (Rtp1-I191AzF), and an acceptor fluorophore attached to the initiation tail of the substrate (Fig. 2D, Table S4). To kinetically isolate tail-insertion from subsequent steps of substrate processing, we treated the proteasome with oPA, thereby preventing progression past the ubiquitin-modified lysine and trapping the proteasome with the inserted substrate in a high-FRET state. After stopped-flow mixing of acceptor-labeled substrate and donor-labeled, oPA-treated proteasome, we observed a rapid increase in FRET as measured by the quenching of donor fluorescence. This was accompanied by a reciprocal increase in acceptor fluorescence, confirming that the observed signal change was caused by an energy transfer between the donor and acceptor (Fig. S4). Fitting of the FRET change revealed a time constant of 1.6 s (Fig. 2D). Since ubiquitin binding to the proteasome has previously been found to occur significantly faster(*41*), we can conclude that the 1.6 s time constant is largely determined by substrate-tail insertion after ubiquitin interaction with a proteasomal receptor. The tail insertion occurred on a time scale similar to the change in substrate anisotropy seen upon mixing with Rpn11^AXA^-containing proteasome (Fig. S4). No signal change was observed when mixing proteasome with substrate lacking a ubiquitin chain (Fig. 2D), and we also confirmed that the fluorophore on the unstructured initiation region had no major effects on the rate of substrate degradation (Table S4).

**Fig. 3.**
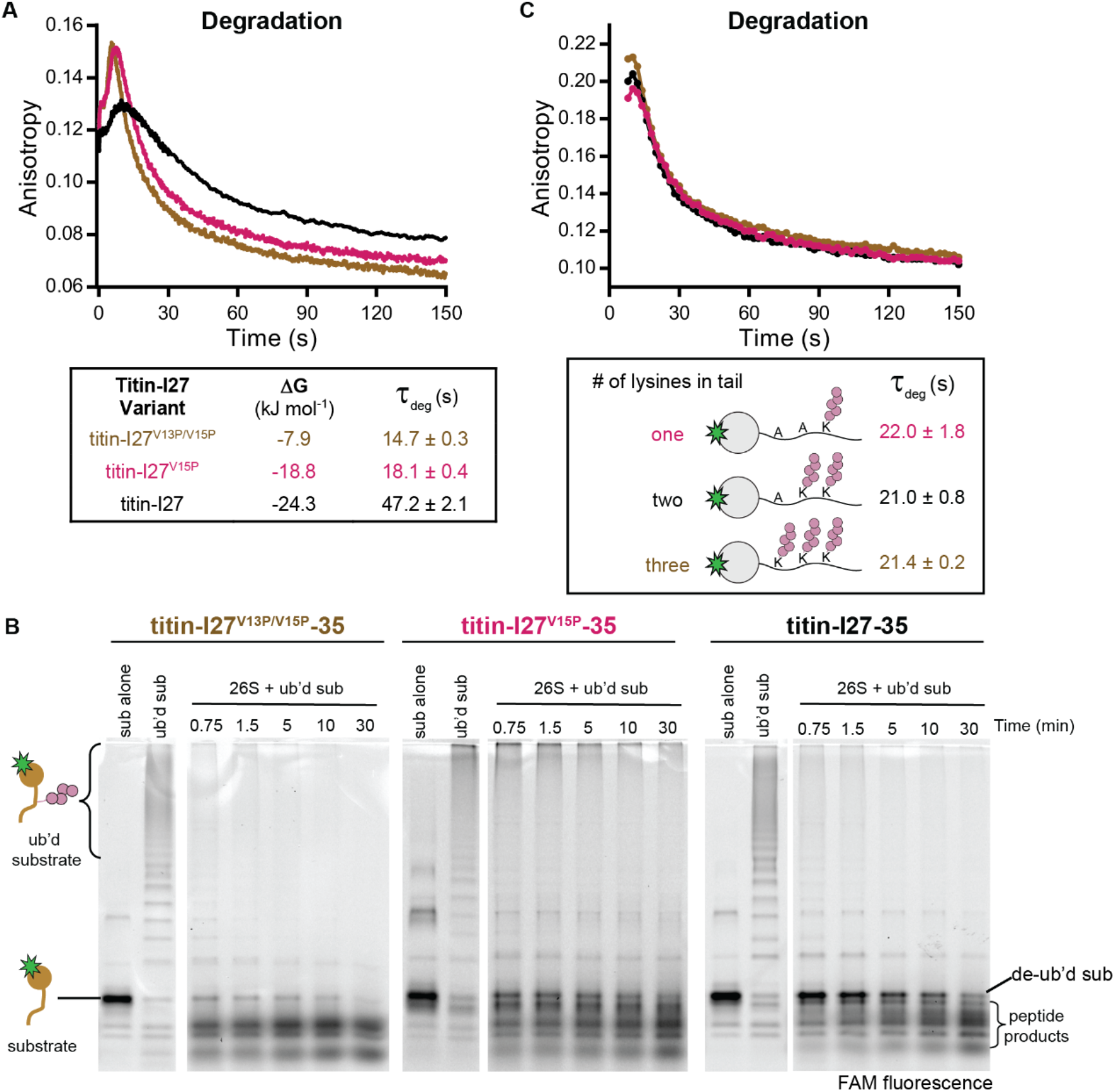
Increasing substrate stability slows degradation, but multiple ubiquitin chains can be rapidly removed. **(A)** Single-turnover degradations of substrates with mutations in the titin-127 folded domain tracked by anisotropy of N-terminally attached 5-FAM in a stopped-flow fluorimeter. The stabilities of titin-I27^v13P/v15P^-35, titin-I27^v15P^-35, and titin-I27-35 were determined by denaturant-induced equilibrium unfolding (see Fig. S6). **(B)** Single-turnover degradations of the same substrates analyzed in A, but tracked by SDS-PAGE and visualized by 5-FAM fluorescence. **(C)**Single-turnover degradations of substrates with up to 3 lysine-attached ubiquitin chains in the unstructured tail tracked by anisotropy of 5-FAM. All curves are representative traces, and time constants are derived from averaging fits of independent experiments, shown with s.d.(*N* = 3).

**Fig. 4.**
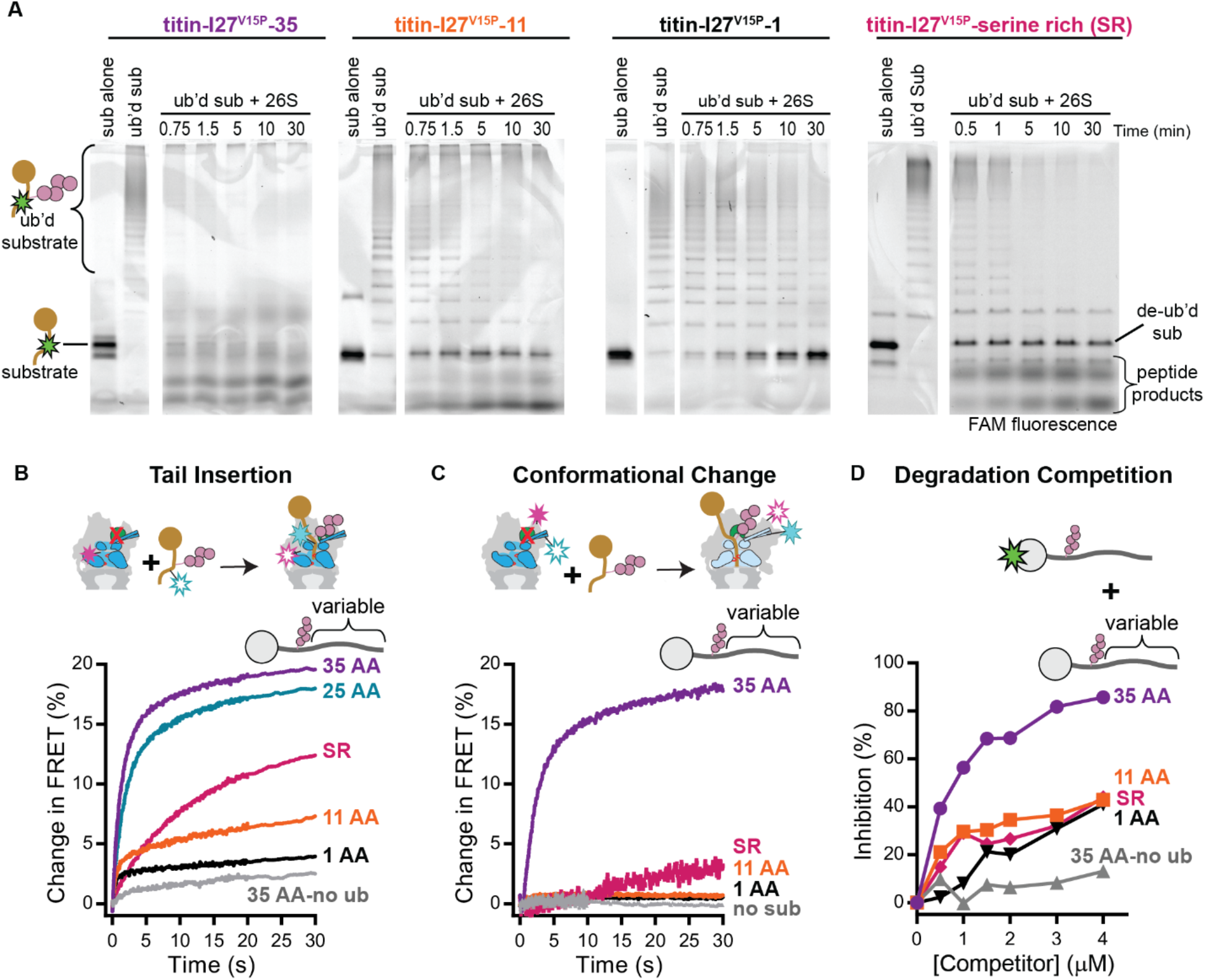
Substrates with poor initiation regions do not stably engage with the proteasome. **(A)** Single-turnover degradations of substrates with varied unstructured initiation regions analyzed by SDS-PAGE and visualized by 5-FAM fluorescence. **(B)** Tail insertion kinetics of substrate variants tracked by FRET between the substrate and the proteasome as in Fig. 2D. The corresponding time constants are listed in Table 1. **(C)** Conformational change of the proteasome after substrate addition, tracked by FRET between the base and the lid as in Fig. 2E. **(D)** Competitive inhibition of titin-I27^v15P^-35 degradation by substrates with varied tails (same substrates as in C). The multiple-turnover degradation of 5-FAM-titin-I27^v15P^-35 at 500 nM was measured by anisotropy in the presence of varying concentrations of unlabeled competitors. Shown is the percent inhibition as derived from the initial rates of degradation in the presence and absence of competitor. All curves are representative traces (N≥ 3).

**Table 1:** Fast phase kinetics and amplitudes of substrate tail insertion as seen in Fig. 4B. Shown are the mean and s.d. for *N* ≥3.

| Substrate | $\tau_{\text{app}}$ (s) | $k_{\text{app}}$ ($1/\tau_{\text{app}}$, $\text{s}^{-1}$) | Amplitude<br>(Normalized to 35 AA tail) |
| --- | --- | --- | --- |
| **titin-I27<sup>V15P</sup>-35** | $1.61 \pm 0.32$ | $0.62 \pm 0.12$ | $1 \pm 0.03$ |
| **titin-I27<sup>V15P</sup>-25**<br>**(serine rich)** | $7.15 \pm 1.00$ | $0.14 \pm 0.02$ | $0.51 \pm 0.04$ |
| **titin-I27<sup>V15P</sup>-25** | $1.73 \pm 0.02$ | $0.58 \pm 0.007$ | $0.87 \pm 0.05$ |
| **titin-I27<sup>V15P</sup>-11** | $0.7 \pm 0.09$ | $1.4 \pm 0.18$ | $0.24 \pm 0.01$ |
| **titin-I27<sup>V15P</sup>-1** | $0.46 \pm 0.03$ | $2.2 \pm 0.14$ | $0.16 \pm 0.001$ |

Tail insertion could be prevented by the pre-incubation of the proteasome with ATPγS (Fig. 2D). In the ATPγS-induced s3 state of the proteasome, Rpn11 is relocated to directly above the central pore and therefore expected to sterically interfere with the insertion of a substrate’s initiation region. The lack of a FRET signal when mixing substrate with ATPγS-bound proteasomes hence confirms that initial binding to a ubiquitin receptors does not lead to FRET and that substrate engagement by the AAA+ motor requires the s1 conformation of the proteasome.

We next measured the kinetics of conformational switching after substrate addition to Rpn11-inhibited proteasomes, using the above-mentioned FRET-based assay that tracks the relative distance between the base and lid sub-complexes (Fig. 2E, Table S5). Because binding of unanchored ubiquitin chains alone had no effect on the conformational state of the proteasome, we hypothesized that engagement of the substrate’s unstructured region with the pore loops of the AAA+ motor is required to trigger the switch. Indeed, upon mixing proteasome with substrate, the acceptor fluorescence increased with a time constant of 2.2 s, closely tracking with the tail insertion event. A reciprocal signal change was observed in the donor channel, confirming the underlying FRET (Fig. S4). Examination of early time points revealed that the FRET signal for substrate tail insertion increased immediately after mixing, whereas the signal for the conformational switch showed a 0.4 s delay (Fig. 2F). This delay indicates that pore-loop contacts with a substrate must occur first, and that this interaction then leads to a rapid switch of the proteasome from the s1 to a substrate-engaged, s3-like state. This model is also supported by our previous studies showing that the pore-loops of Rpt subunits at the top of the s1-state spiral staircase, which are thought to make first contact with a substrate polypeptide, are particularly important for degradation(*18*).

After engagement of a substrate, the proteasome must remove attached ubiquitin chains before proceeding to unfold and translocate the rest of the polypeptide. Therefore, a final FRET-based assay was designed to selectively monitor the deubiquitination event, this time by tracking the energy transfer between donor-labeled ubiquitin and an acceptor fluorophore attached to the substrate adjacent to the ubiquitinated lysine. Co-translocational ubiquitin removal from the substrate caused a loss in acceptor fluorescence, while pre-incubation of proteasomes with ATPγS abolished the signal change (Fig. 2G, Fig. S4D). Stopped-flow measurements of the FRET signal upon mixing substrate with saturating amounts of wild-type proteasome in the presence of ATP revealed that deubiquitination occurs with a total time constant of 6.8 s (Table S6), which reflects the sum of the time required for binding, tail insertion, and deubiquitination. Mixing of the substrate with Rpn11^AXA^-containing proteasome also led to a signal change, but with more than 50% reduced amplitude. This deubiquitination-independent change in FRET is likely caused by a change in fluorophore environment when ubiquitin binds to the catalytically-dead Rpn11 (Fig. S5).

**Fig. 5.**
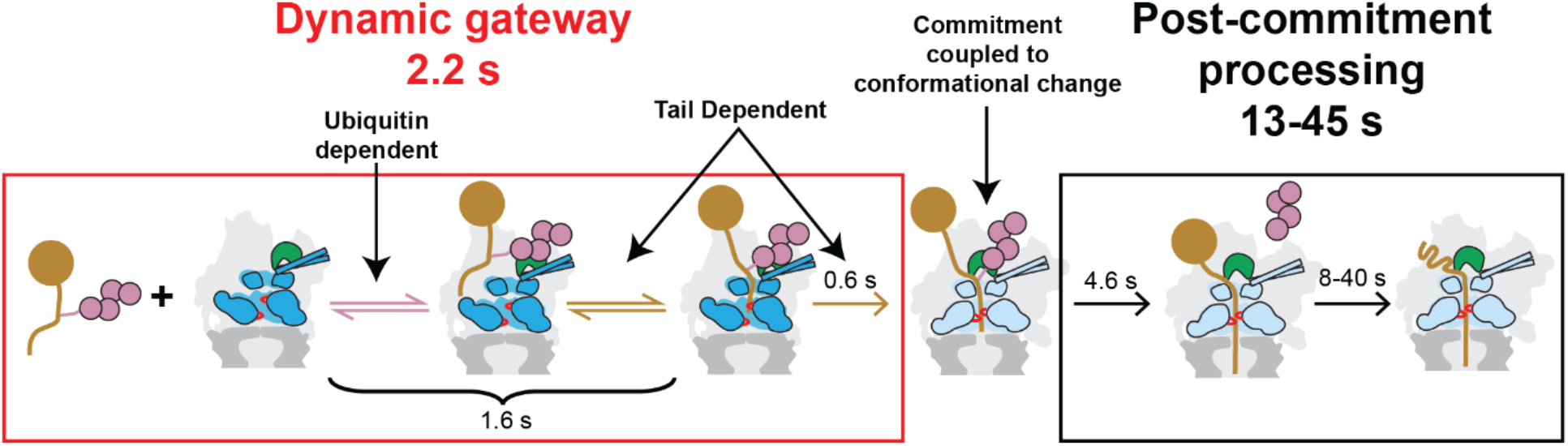
Model for kinetic proofreading by the proteasome. Ubiquitin binding and tail insertion constitute a dynamic gateway to proteasomal substrate processing, with both fast on and off rates. If the substrate has the necessary requirements for degradation, engagement with the AAA+ motor reduces k_off_ for tail insertion and accelerates the conformational switch of the proteasome, thereby committing substrates to translocation-coupled deubiquitination, rate-limiting unfolding, and degradation.

By comparing the various time constants described above, we can derive the first complete kinetic model of proteasomal substrate processing and estimate the time required for each sub-step following substrate binding (Fig. 2H). Initial tail insertion proceeds with a time constant of 1.6 s, followed by a rapid conformational switch after 0.4 s. The removal of the ubiquitin chain then happens within ~ 5 s, and mechanical substrate unfolding, translocation, and cleavage take an additional 11 s.

## The proteasome can quickly remove multiple ubiquitin chains, but is slowed down by stable domains

Our kinetic characterization of proteasomal degradation revealed that most of the substrate processing time is spent on deubiquitination and unfolding or translocation of the substrate. To further probe the relative contributions of these steps to overall degradation time, we modified our model substrate, either by changing the stability of the folded domain (Fig. 3A,B Fig. S6) or by increasing the number of ubiquitin chains that need to be removed during degradation (Fig. 3C, Fig. S7).

A series of substrate variants was constructed based on previously reported mutations that change the thermodynamic stability of the titin-∆27 domain(*42*). The experiments reported in Fig. 2 were performed with titin-I27^V15P^-35, which has a thermodynamic stability of ΔG = −19 kJ mol^−1^ (Fig. S6). In addition, we measured the degradation rates for a further destabilized variant titin-I27^V13P/15P−^35 (∆G = −8 kJ mol^−1^) and for the wild-type version, titin-I27-35 (ΔG = −24 kJ mol^−1^). Increasing the stability of the folded titin domain significantly increased the overall time required for degradation, as seen for other substrates(*43-45*). Further analysis by SDS-PAGE showed that the most stable titin variant is rapidly deubiquitinated and then slowly unfolded and degraded (Fig. 3B), indicating that the substrate remains stably engaged after ubiquitin-chain removal and while the proteasome attempts to overcome the tough unfolding barrier. Thus, the unfolding event itself, as opposed to tail insertion, deubiquitination, or translocation, is the rate-limiting step of degradation for this substrate. The less stable titin variants exhibited no accumulation of the deubiquitinated species on the proteasome, indicating that unfolding and deubiquitination occur on similar time scales for these substrates. Interestingly, the stability of the titin variants correlated with their ability to stimulate proteasomal ATP hydrolysis. The most stable substrate (titin-I27-35) showed the strongest stimulatory effect (Table S7), most likely because its slow unfolding caused the proteasome to spend the least time in the non-stimulated, substrate-free state during degradation under saturating conditions.

The least stable titin-I27^V13P/15P^ variant was degraded with a total time constant of 15 s. Considering that all initial processing steps, including deubiquitination, take 6.8 seconds, and assuming rapid subsequent unfolding of the destabilized titin domain, we can estimate a minimum translocation rate of 15 residues s^−1^ for the 116 amino acids that remain to be threaded after unfolding.

To test the effects of additional ubiquitin chains on substrate degradation, we added a second or third lysine into the unstructured tail of our model substrate. The modification of all three lysine was confirmed by SDS-PAGE after ubiquitination with methyl-ubiquitin (Fig. S7A,B) or by treating the poly-ubiquitinated substrate with the deubiquitinase AMSH (Fig. S7C), which trims K63-linked chains, but does not remove the substrate-attached ubiquitin moiety(*46*). Michaelis-Menten analyses showed that substrates with one, two, or three ubiquitin chains have similar binding affinities and degradation rates (Fig. S2). Importantly, all three substrates also showed no difference in degradation rate under single-turnover conditions (Fig. 3C), suggesting that while the removal of the first ubiquitin chain takes 5 seconds, subsequent chains must be removed much more quickly. One model for this change in deubiquitination rate comes from our previous findings that the rate-limiting step of isopeptide cleavage by Rpn11 is a ubiquitin-induced conformational switch of its Insert-1 region(*21*). We propose that after Rpn11 removes the first ubiquitin chain, it does not immediately switch back to the inactive conformation, but instead is poised to bind and promote fast cleavage of nearby ubiquitin chains that rapidly approach Rpn11 as the substrate is translocated into the central pore. Many essential proteasome substrates, such as the cell-cycle regulators ubiquitinated by the anaphase-promoting complex (APC), are known to be modified with multiple ubiquitin chains(*47, 48*). Our data suggest that removal of ubiquitin chains is not rate-limiting for the degradation of these substrates.

## Substrates with poor initiation regions fail to engage with the proteasome

It has been established that substrates with short (< 25 residues) or low-complexity initiation regions are not efficiently degraded *in vitro* and have extended *in vivo* half-lives(*4-7, 49*). To investigate the mechanisms behind this defect, we modified the initiation tail of our model substrate, either by varying the length of the terminal unstructured region following the single ubiquitinated lysine or by replacing it with a low complexity, serine-rich sequence previously shown to severely impair degradation(*5, 49*).

Tracking of proteasomal processing by SDS-PAGE revealed that substrates with 35 or 25 AA tail were completely degraded into peptides, whereas the truncated tail variants with 11 or 1 AA were primarily deubiquitinated and released (Fig 4A, Fig. S8, Fig. S9). Deubiquitination and release of the short-tail substrates could be monitored by the decrease in anisotropy, though the magnitude of signal change was of course smaller than for complete proteolysis into peptides (Fig. S10). Using this anisotropy readout in multiple turnover titrations revealed that the short tails cause significant K_M_ defects for the titin-I27^V15P^-11 (KM = 1.2 μΜ) and titin-I27^V15P^-1 (K_M_ = 2.7 μΜ) substrates, compared to titin-I27^V15P^-35 (K_M_ = 0.2 μΜ) (Fig. S2, S11).

We then analyzed the kinetics of tail insertion through the increase of FRET between donor labeled, Rpn11-inhibited proteasome and acceptor-labeled substrate, as in Fig. 2D (Fig. 4B, Table 1). We found that shorter tails appear to enter the central pore more rapidly, yet their FRET amplitude was severely reduced, with 11 and 1 AA tails showing only 25% and 16%, respectively, of the signal change detected for the 35-residue tail. These low amplitudes are most likely a consequence of the truncated tails being too short to reach from the narrow N-ring entrance to the pore loops, thus preventing a stable engagement with the AAA+ motor. After initial rapid insertion, these tails may therefore quickly escape again from the central channel, which is consistent with the observed increase in the apparent rate constants for short tail insertion, k_app_, that reflects the sum of both, on and off rates (Table 1). The elevated off rates lead to an increase in K_M_ for the processing of short-tailed substrates under multiple-turnover conditions, even though their ubiquitin targeting signal is unchanged (Fig. S11).

Though short tails insert, at least briefly, into the pore of the motor, we found that substrates with those truncated initiation regions fail to trigger the conformational change of the proteasome (Fig. 4C). In contrast to the small amplitude and fast kinetics we observed for the tail-insertion traces, the conformational change traces for the truncated tails exhibited almost no amplitude. The inability of short-tailed substrates to induce a conformational change thus supports our model that interactions between the substrate polypeptide and the motor’s pore loops drive the switch in proteasome conformation.

Despite their failure to engage and trigger proteasome conformational switching, short-tailed substrates are slowly deubiquitinated (Fig. 4A). The loss of anisotropy during the multiple-turnover processing of the 1 AA tail substrate (Fig. S11) reveals a time constant of 45 s for this engagement-independent deubiquitination, which is significantly slower than the 6.8 s observed for translocation-coupled deubiquitination of the long-tailed substrate (Fig. 2G). The almost 7-fold difference in deubiquitination rate agrees well with the previously revealed acceleration of Rpn11-mediated ubiquitin cleavage by mechanical substrate translocation into the AAA+ motor (*21*). Since short-tailed substrates fail to induce the conformational switch of the proteasome, their slow deubiquitination likely occurs while the proteasome is in the s1 state or during brief, spontaneous sampling of s3-like conformation. Despite not being coupled to translocation, this engagement-independent ubiquitin cleavage still results in the accumulation of only fully deubiquitinated substrate (Fig. 4A), as expected if steric hindrance by the subjacent N-ring of the AAA+ motor forces Rpn11 to cleave at the base of the ubiquitin chain, rather than between ubiquitin moieties.

Like the short tails, the low complexity serine-rich tail also led to a significant K_M_ defect in multiple-turnover processing and exhibited a decreased FRET amplitude for tail insertion (Fig. S11, Fig. 4B). In contrast to the short tails, however, it also showed slower tail-insertion kinetics, with a k_app_ four times lower than that of a complex tail of the same length. Thus, in addition to having an elevated ratio of k_off_/k_on_ (and therefore K_M_), the on-rate of the serine-rich tail is decreased. The conformational change induced by this substrate was also slow and of a low magnitude, with the defects again being amplified compared to those measured in tail insertion. Both the tail-insertion and conformational change assays rely on a stalled state, with the substrate’s initiation region stably engaged by the AAA+ motor. The small FRET-signal amplitudes for the serine-rich tail thus indicate a rapid substrate escape from the proteasome pore and a consequent low probability of commitment, explaining the observed slow degradation and frequent release (Fig. 4A). Low-complexity regions thus hinder substrate processing in several ways: by slowing the onset of tail insertion, inhibiting the conformational change, and likely by reducing the motor grip required for unfolding and translocation.

How are proteasomes *in vivo* able to rapidly turn over high priority substrates, such as cell-cycle regulators, while ubiquitinated proteins with poor initiation regions and therefore slow processing kinetics compete for proteasomal ubiquitin receptors? To test this scenario *in vitro,* we performed competition experiments, in which the degradation of a fluorescently labeled, ubiquitinated titin-I27^V15P^-35 substrate was tracked in the presence of unlabeled ubiquitinated substrates with varying tail architectures (Fig. 4D). While the unlabeled substrate with a 35 residue tail effectively inhibited degradation of its labeled counterpart, substrates with short or serine-rich initiation regions were poor competitors, even at concentrations well above their measured K_M_ values. These results can be explained by the high off rates and poor engagement of these tails with the ATPase motor. Their inability to compete also suggests that the ubiquitin-chain interactions with receptors on the proteasome is short lived. The translocation-independent deubiquitination we observed for substrates with poor initiation regions (Fig. 4A) must happen during brief and presumably repetitive binding events. The proteasome thus has intrinsic mechanisms for removing poor substrates from the pool of ubiquitinated proteins, while still allowing the preferred and therefore rapid degradation of appropriate substrates. Accessory factors such as Ubp6 and Hul5, both proposed to be important for substrate triage, may act in concert with the proteasome’s intrinsic mechanisms described here(*50*).

## Conclusions

An intricate system of recognition and processing steps allows the 26S proteasome to select only appropriate substrates that contain both the correct ubiquitin-targeting signal and an unstructured initiation region, while maintaining the high promiscuity necessary to degrade hundreds of cellular proteins with diverse characteristics. Using FRET-based assays to measure the kinetics of individual substrate-processing events, we found that insertion and engagement of the unstructured initiation tail is fast, whereas most of the degradation time is spent on mechanical unfolding and translocation. We show that the proteasome rapidly transitions from the s1 state to a substrate-processing, s3-like state immediately after tail insertion, suggesting that substrate interactions with the pore loops of the AAA+ motor drive this global conformational switch. In the s1 state, the entrance to the central pore is well accessible for the substrate’s flexible initiation region, and the ATPase domains of Rpt1-6 are arranged in a steep spiral staircase that likely helps substrate entry(*18, 28, 31*). The transition to the s3-like processing state, right after substrate engagement with the translocation machinery, is ideally timed to facilitate all subsequent processing steps, as it brings the ATPase ring into a more planar, translocation-competent conformation and moves Rpn11 to directly above the central pore to enable efficient coupling of translocation and deubiquitination(*21, 30*). Premature transition to this state, however, prevents tail insertion and engagement, which highlights the importance of this intricate coordination between substrate processing steps and conformational switching of the proteasome.

We identified mechanical unfolding as the rate-determining step for degradation of our model proteins and likely most proteasomal substrates in general. Consistent with previous studies, we found that increasing the thermodynamic stability of a substrate’s structured domain extends the time required for degradation. In contrast, the presence of additional ubiquitin chains had no effect on the overall degradation rate. We propose that after the first ubiquitin-cleavage, which takes about 5 s, subsequent deubiquitination events can be significantly accelerated, because Rpn11 remains in its active state at least long enough to remove closely spaced ubiquitin chains from a translocating polypeptide. Multiple ubiquitin chains can thus confer additional affinity for the proteasome without reducing the degradation rate, and it is not surprising that high-priority substrates, such as those ubiquitinated by the APC, contain multiple, closely spaced ubiquitin modifications(*47, 48*). While the removal of ubiquitin chains may not be limiting, their orientation relative to a substrate’s unstructured initiation region has been found to potentially affect the rate of degradation(*6, 41*). Future studies, utilizing the specific assays presented here, will have to address whether those differences in turnover rates are due to altered engagement kinetics or differential rates of deubiquitination depending on the spacing of ubiquitin chains relative to each other or a folded domain.

When investigating the requirements for substrate recognition and engagement, we found that ubiquitinated proteins with initiation regions of insufficient length or complexity were slowly deubiquitinated and released, rather than degraded, which explains previous findings of poor *in vitro* degradation or extended *in vivo* half-lives for those substrates(*7, 9*). Short or low-complexity initiation regions still rapidly enter the central channel of the proteasome, but cannot stably engage with the AAA+ pore loops and quickly escape again, leading to a significant increase of the substrate’s K_M_. Remodeling factors such as Cdc48/p97 may thus play a critical role in preparing substrates for degradation by exposing unstructured initiation regions for efficient proteasomal engagement(*51, 52*).

We propose that the initial ubiquitin-binding and tail-insertion events represent a dynamic gateway that is characterized by both, fast on and off rates, and controls the selection of appropriate substrates for proteasomal degradation (Fig. 5). If a substrate has the necessary requirements for processing, then tail engagement with the AAA+ motor reduces the off rate and induces the proteasome conformational switch that commits the substrate to unfolding, translocation-coupled deubiquitination, and degradation. As a consequence of this rapid ubiquitin sampling by the proteasome, binding of poor substrates does not considerably inhibit the degradation of higher-priority substrates with proper initiation regions, as seen in our *in vitro* competition experiments with short and long tailed proteins. This mechanism is thus similar to the kinetic proofreading mechanism used by the ribosome to recognize cognate amino acyl-tRNAs, in which rapid initial binding interactions are followed by an irreversible committed step(*53*). In addition to this proofreading, the slow, translocation-independent deubiquitination of engagement-incompetent proteins at the proteasome removes them from the pool of potential substrates. Our studies thus provide exciting new insights into how the proteasome prioritizes its substrates in the cell and coordinates its processing steps and conformational changes to ensure efficient protein degradation.

## Acknowledgements

We thank the members of the Martin lab for numerous helpful discussions. Funding: J.A.M.B. and C.B. acknowledge support from NSF Graduate Research Fellowships. This research was funded in part by the US National Institutes of Health (R01-GM094497 to A.M.), the US National Science Foundation CAREER Program (NSF-MCB-1150288 to A.M.), and the Howard Hughes Medical Institute (K.C.D., and A.M.).

## Author contributions

J.A.M.B. and A.M. designed experiments, J.A.M.B. performed biochemical experiments, C.B. helped with experimental design, and K.C.D. assisted with the preparation of materials. All authors contributed to manuscript preparation.

### Data and materials availability

All data are available in the main text or the supplementary materials. Supplementary Materials

## Materials and Methods

All plasmids used for the purification of recombinant proteins are listed in Table S1.

### Purification of 20S core

*S. cerevisiae* 20S core was purified from the Pre1-3xFLAG yeast strain yAM14 (RJD1144, from Ray Deshaies,(54)) as previously described(30). Briefly, yAM14 yeast were lysed and resuspended in lysis buffer (60 mM HEPES, pH 7.6, 500 mM NaCl, 1 mM EDTA, 0.2% NP-40). 20S core was immobilized on M2 anti-FLAG resin (Sigma), washed with buffer containing 1 Μ NaCl to remove bound regulatory particle, eluted from the resin using 3xFLAG peptide, and further purified by size-exclusion chromatography with a Superose 6 Increase 10/300 column (GE) equilibrated in GF buffer (30 mM HEPES, pH 7.6, 50 mM NaCl, 50 mM KCl, 10 mM MgCl_2_, 5% glycerol) with 0.5 mM TCEP. Peak fractions were concentrated in a 100K MWCO Amicon Ultra concentrator (Millipore), aliquoted, flash frozen in liquid N_2_, and stored at −80 °C. The concentration was determined by absorbance at 280 nm.

### Purification of wild-type base

Recombinant yeast base was purified after heterologous expression in *E. coli* using a slight modification of previously published methods *(18). E. coli* BL21 Star(DE3) (Invitrogen) was co-transformed with the plasmids pAM81, pAM82, and pAM83 (derived from pETDuet, pCOLADuet, and pACYCDuet, respectively), coding for Rpt1 - Rpt6, Rpn1, Rpn2, Rpn13, the four base-assembly chaperones Rpn14, Hsm3, Nas2, and Nas6, and rare tRNAs. Bacteria were grown in 3 L of terrific broth (Novagen) to an OD_600_ = 1.0, then induced with 0.5 mM IPTG at 30 °C for 5 hours, followed by an overnight induction at 16 °C. The cells were pelleted and resuspended in NiA buffer (60 mM HEPES, pH 7.6, 100 mM NaCl, 100 mM KCl, 10 mM MgCl_2_, 5% glycerol, and 20 mM imidazole) supplemented with 2 mg ml^−1^ lysozyme, benzonase (Novagen), and protease inhibitors (aprotinin, pepstatin, leupeptin and PMSF). Cells were lysed by sonication, the lysate was clarified by centrifugation, and the base subcomplex was purified by a three-step procedure, with 0.5 mM ATP present in all buffers. First, the His-Rpt3 containing complexes were purified using a 5 mL HisTrap FF crude (GE), then fully assembled complexes containing Flag-Rpt1 were selected for by using anti-Flag M2 affinity resin (Sigma), and finally, the complex was further purified using size-exclusion chromatography with a Superose 6 increase 10/300 column (GE) equilibrated in GF buffer with 0.5 mM ATP and 0.5 mM TCEP. The concentration of the base was determined by Bradford assay using bovine serum albumin (Sigma) as a standard.

### Purification and labeling of unnatural amino acid-containing base

The base sub-complex containing 4-azido-L-phenylalanine (AzF) was expressed, purified, and labeled using a similar procedure with the following modifications: In addition to pAM81, pMA83, and an amber-codon containing variant of pAM82, a fourth plasmid (pAM87) containing the AzF tRNA synthetase/tRNA pair was used to co-transform Bl21 Star(DE3) *E. coli.* This plasmid was constructed by replacing the IPTG-inducible synthetase in the pULTRA-CNF construct designed by the Schultz lab with the AzFRS.2.t1 synthetase evolved by the Isaacs lab(*34, 35*). The pEvol-pAzFRS.2.t1 was a gift from Farren Isaacs (Addgene plasmid # 73546) and the pULTRA-CNF was a gift from Peter Schultz (Addgene plasmid # 48215). The cells were grown in 3 L of dYT media to an OD_600_ = 0.6, then pelleted, and resuspended in 0.5 L of terrific broth (Novagen) with added 2mM AzF (Amatek Chemical). The cells were shaken for 30 minutes at 30 °C to allow for uptake of the AzF, followed by induction with 0.5 mM IPTG for 5 hours at 30 °C and then 16 °C overnight. Purification followed the same protocol as for AzF-free base, except that after elution from the anti-Flag resin, the base was incubated with 150 μM 5,5’-dithiobis-(2-nitrobenzoic acid) (DTNB) for 10 minutes at room temperature to temporarily block any exposed cysteines. The base was then cooled back down to 4 °C and labeled overnight with 300 μM dibenzocyclooctyne (DBCO) conjugated fluorophore (Click Chemistry Tools). The reaction was quenched using 1 mM free AzF, followed by 2 mM dithiothreitol (DTT), and labeled base was then purified using size-exclusion chromatography as described above. Labeling efficiencies were estimated by comparing the absorbance-based concentration of fluorophore with the base concentration calculated by the Bradford assay.

### Purification of lid

Recombinant yeast lid was also purified using a three–step procedure. The plasmids pAM80, pAM85, and pAM86, coding for Rpn3, Rpn5 - Rpn9, Rpn11, Rpn12, Sem1, and rare tRNAs, were modified from previously published plasmids to replace the affinity tags*(31)*. Human rhinovirus (HRV) protease-cleavable maltose binding protein was inserted at the N-terminus of Rpn6, and the N-terminus of Rpn12 was modified with a HRV-cleavable 6xHis tag. *E. coli* BL21 star(DE3) was co-transformed with these plasmids and then grown, induced, and lysed as described above for the base. The fully assembled lid was purified using a HisTrap and amylose resin (NEB), cleaved with HRV-protease, and purified on a Superose 6 Increase size-exclusion column equilibrated in GF buffer with 0.5 mM TCEP. The concentration of the lid was determined by absorbance at 280 nm. Unnatural amino acid-containing lid was expressed and purified in a fashion similar to AzF-containing base, using the same DTNB blocking and DBCO-fluorophore labeling procedures after elution from the amylose resin.

### Purification of substrates

All substrates were purified using the IMPACT system (NEB). The substrate was inserted into a T7-inducible plasmid derived from pET28a (Novagen) upstream of the intein and chitin binding domain (CBD) taken from pTXB1 (NEB). *E. coli* BL21 star(DE3) was transformed with this plasmid, grown in 3 L of dYT to an OD_600_ = 0.6, and induced with 0.5 mM IPTG for 3 hours at 30 °C. The cells were then resuspended in chitin buffer (60 mM HEPES, pH 7.6, 150 mM NaCl, 1 mM EDTA, 5% glycerol) with protease inhibitors (AEBSF, aprotinin, leupeptin, and pepstatin), and benzonase (Novagen). The cells were bound in batch to 10 mL of chitin resin (NEB) for 1 hour at 4 °C, which was then washed with 100 mL of chitin buffer supplemented with an additional 500 mM NaCl, 0.2% Triton X-100, and protease inhibitors. The substrate was cleaved from the column by overnight incubation at 4 °C in a buffer containing 60 mM HEPES, pH 8.5, 150 mM NaCl, 1 mM EDTA, 5% glycerol, and 50 mM DTT. The substrate was collected from the column, run over 3 mL of fresh chitin resin to remove any uncleaved substrate, and then purified on a S75 16/60 size-exclusion chromatography column (GE) in GF buffer with 0.5 mM TCEP. The concentration of substrate was determined by the absorbance at 280 nm.

### Labeling of substrates

Fluorophores were covalently attached to a single engineered cysteine in the substrate by maleimide chemistry. While there are two additional cysteines in the titin-I27 domain used, they are not solvent exposed. Where noted, the N-terminus of the substrate was labeled by sortase A (SrtA-mediated ligation of a peptide (55). For the sortase modification, peptide (HHHHHHLPETGG) was purchased with 5-FAM conjugated to its N-terminus (Biomatik), and then ligated to 100 μM substrate using 20 μM recombinant 6xHis-SrtA in GF buffer supplemented with 10 mM CaCl_2_, and 0.5 mM peptide. The labeled substrate was selected for by binding to a 1 mL HisTrap HP (GE), followed by size-exclusion chromatography on a Superdex 75 10/300 column (GE) equilibrated with GF buffer. The concentration of substrate was determined by quantifying the absorbance of the attached fluorophore (5-FAM = 492 nm, Cy5 = 646 nm).

### Purification and labeling of ubiquitin

In order to label ubiquitin, a variant was expressed with an N-terminal cysteine inserted before the first methionine (MC-ubiquitin). This variant was expressed and purified as described previously (21). Briefly, Rosetta II (DE3) pLysS *E. coli* cells were transformed with an IPTG-inducible expression plasmid (pET28a) containing MC-ubiquitin. The cells were grown in terrific broth (Novagen) at 37 °C until the OD_600_ = 1.5, and ubiquitin expression was induced with 0.5 mM IPTG overnight at 18 °C. After expression, the cells were resuspended in lysis buffer (50 mM Tris-HCl, pH 7.6) containing 2 mg mL^−1^ lysozyme, benzonase, and protease inhibitors (aprotinin, pepstatin, leupeptin and PMSF). The cells were lysed by sonication, and contaminating proteins were precipitated by the addition of 60% perchloric acid to a final concentration of 0.5%. The soluble fraction containing ubiquitin was dialyzed overnight into 50 mM Na-acetate, pH 4.5, and purified by cation exchange on a 5 mL HiTrap SP FF column (GE) using a gradient of 0 - 0.5 M NaCl in 50 mM Na-acetate, pH 4.5. Peak fractions were concentrated and further purified over a Superdex S75 16/60 column in storage buffer (20 mM Tris-HCl, pH 7.6, 150 mM NaCl, 1 mM DTT). Ubiquitin was labeled by dialyzing into labeling buffer (30 mM HEPES, pH 7.2, 150 mM NaCl, 1 mM EDTA) and reacted with an excess of maleimide-fluorophore. The labeled ubiquitin was purified on a Superose 75 16/60 size exclusion column equilibrated in GF buffer, and its concentration was measured by the Lowry assay using non-labeled ubiquitin as a standard.

A pET15b plasmid containing linearly fused tetra-ubiquitin with an N-terminal fusion to 6xHis-SUMO was generated by gene synthesis (Genscript). The protein was expressed in Rosetta II (DE3) pLysS *E. coli,* purified on a HisTrap, cleaved using SENP1 protease, then further purified by cation exchange on an SP FF column (as described above) and size-exclusion chromatography on a Superose 75 16/60 column equilibrated in GF buffer.

### Purification of additional components

The plasmid used to express Usp2 was a gift from Cheryl Arrowsmith (Addgene plasmid # 36894). The plasmid used to express SENP2 protease was a gift from Guy Salvesen (Addgene plasmid # 16357)(*56*). Both proteins were expressed in *E. coli* BL21 star(DE3) and purified using a HisTrap, followed by size-exclusion chromatography on a Superose 75 16/60 column equilibrated in GF buffer. The plasmid used to express mouse E1 (pet28-mE1) was a gift from Jorge Eduardo Azevedo (Addgene plasmid # 32534)(*57*). The plasmid used to express AMSH, pOPINB-AMSH*, was a gift from David Komander (Addgene plasmid # 66712) and was purified as described previously(*46*).

Additional recombinant proteins including Rpn10, ubiquitin, Rsp5, Ubc1, and E1 were purified as previously described(*21*). Ubp6^C188A^ was purified as previously described(*37*).

### Substrate ubiquitination

All substrates contained a PPPY motif in the unstructured region, which allows them to be ubiquitinated with long primarily K63-linked ubiquitin chains *in vitro* using the Rsp5 E3 ligase (*30, 39*). Purified substrates (10 μM) were incubated for 3 hours at 25 °C in GF buffer with E1 (2.5 μM), Ubc1 (2.5 μM), Rsp5 (2.5 μM), ubiquitin (400 μM), and 10 mM ATP. Substrates were ubiquitinated fresh for each day of experiments and evaluated by gel to ensure reproducible ubiquitination.

## Enzymatic assays

All enzymatic assays were performed at 25 °C in GF buffer supplemented with 5 mM ATP, 0.5 mM TCEP, 0.5 mg mL^−1^ BSA, and an ATP regeneration system (0.03 mg mL^−1^ creatine kinase and 16 mM creatine phosphate) unless otherwise noted. Experiments were performed in at least three technical replicates.

### Data processing and analysis

All curve fitting was done using Origin (Originlab, Northampton, MA). The fits were performed by least squares fitting, and evaluated by the distribution of the residuals to ensure there were no significant deviations from zero. The initial rates from substrate degradation or deubiquitination experiments were fit to the Michaelis-Menten equation to determine k_cat_ and K_M_. Exponential curves from single-turnover measurements were fit to either a double exponential increase/decay (y=y0 + A1*e^−t/τ1^ + A2*e^−t/τ2^) or a delayed double exponential increase/decay (y=y0 + A1*e^−(t-t0)/τ1)^ + A_2_*e^−(t-t0)/τ2^), in which the delay was determined by visual inspection of the curves. In general, the channel with the largest signal change was used for fitting and representative traces. Some curves were smoothened by the Savitsky-Golay method for visualization, but all data analysis was done on the raw traces. The plotted curves were normalized to the initial fluorescence value for each trace.

The slow phase of the double exponential fit was not used in the analyses, because it is thought to arise from a small population of partially aggregated or incompletely ubiquitinated substrate. In support of this, the fast phase of the single-turnover degradation kinetics match well with the kcat values measured in multiple turnover. The time constants for both phases and their relative distribution of amplitudes are reported in Tables S3-S6.

### Anisotropy assays

Multiple-turnover degradations were monitored by tracking fluorescence anisotropy of fluorescein in a Synergy NEO2 multimode plate reader (Biotek). Reactions were initiated by mixing 6.5 μL of 2x reconstituted holoenzyme with 6.5 μL of 2x ubiquitinated substrate, then transferring 10 μL of the reaction to a 384-well flat bottom, low volume microplate (Corning). Final concentrations were 25 or 50 nM core, 400 nM base, 600 nM lid, 750 nM Rpn10, and varying concentrations of substrate.

Initial rates of processing were calculated by using the measured anisotropy of ubiquitinated substrate alone and that of substrate fully cleaved by chymotrypsin to normalize the measured change in anisotropy during the reaction.

To determine the Michaelis-Menten constants for titin-I27^V15P^-1-FAM, the initial rate was normalized using substrate that had been fully deubiquitinated by Usp2 rather than fully degraded substrate. Because titin-I27^V15P^-11-FAM and titin-I27^V15P^-serine rich-FAM exhibit both degradation and deubiquitination, the chymotrypsin degraded value was used to normalize the curves, but the k_cat_ is most likely underestimated.

Final concentrations for all single-turnover degradations were 1.25 μM core, 2.5 μM base, 2.5 μM lid, 3 μM Rpn10, and 300 nM substrate. Single-turnover degradation measurements that were performed in a plate reader used a similar procedure as described above, with a deadtime of 8-10 seconds between mixing and the first measurement. Degradation measurements were also performed in an Auto SF120 stopped-flow fluorimeter (Kintek) with two photomultipliers to measure fluorescein anisotropy. The stopped-flow was loaded with 140 μL of 2x reconstituted holoenzyme and 140 μL of 2x substrate, with the same final concentrations as those used in the plate reader. After loading, four blank shots were fired, followed by three measurements. The deadtime of mixing on the instrument is sub-millisecond.

The curves for degradations measured in the stopped-flow and those measured in the plate reader were fit to a delayed exponential.

Single-turnover degradation measurements that were analyzed by gel were performed by mixing 2x reconstituted holoenzyme and 2x substrate, taking 2 μL aliquots at the indicated time, quenching them in 2% SDS-containing buffer, and separating the reaction by SDS-PAGE on 12% NuPAGE gels (Invitrogen). The fluorescence was then measured on a Typhoon FLA 9500 variable mode scanner (GE) using a pixel density of 50 μm per pixel.

### Tail insertion assays

Tail insertion was measured by tracking Förster resonance energy transfer (FRET) between Cy5-labeled base and Cy3-labeled substrate, while enforcing single-turnover conditions using 1,10-phenanthroline (oPA), an inhibitor of Rpn11 that stalls the proteasome at the site of ubiquitin linkage *(21*). Base containing Rpt1-I191AzF was labeled with Sulfo-Cy5-DBCO (Click Chemistry Tools) as described above. A 30 mM oPA stock was made in GF buffer. Reactions were performed in the Auto SF120 using 2x reconstituted holoenzyme and 2x Sulfo-Cy3 labeled substrate. Final concentrations were 100 nM Cy5-Rpt1 base, 400 nM core, 600 nM lid, 750 nM Rpn10, 3 mM oPA, and 3 μM substrate. An excitation wavelength of 550 nm was used, and the Cy3 and Cy5 emission channels were monitored simultaneously.

The kinetics of tail insertion were determined by fitting the quenching of the Cy3 signal. The amplitudes listed in Table 1 were calculated by normalizing the amplitudes of the fast-phases to the initial fluorescence, and then comparing those to the normalized amplitude of titin-I27^V15P^-35-Cy5. All measurements used in this calculation were from traces collected on the same day.

### Conformational change assays

The conformational state of the proteasome was monitored using FRET between Cy3-labeled lid and Cy5-labeled base. Lid containing Rpn9S111AzF was labeled with Sulfo-Cy3-DBCO, and base containing Rpt5Q49AzF was labeled with Sulfo-Cy5-DBCO as described above. Steady-state measurements were performed by mixing 2x reconstituted holoenzyme (150 nM Cy5-Rpt5 base, 400 nM core, 500 nM Cy3-Rpn9 lid, and 750 nM Rpn10) with 2x substrate or other additive as indicated. Final concentrations of the reconstituted holoenzyme were 150 nM Cy5-Rpt5 base, 400 nM core, 500 nM Cy3-Rpn9 lid, and 750 nM Rpn10. Final substrate concentration was 3 μM. Final oPA concentration was 3 mM. Catalytically dead Ubp6(C188A) was added at 250 nM, and linear tetra-ubiquitin was added at 100 μM. ATPγS was added at 5 mM in place of the ATP regeneration system. The reactions were then transferred to a 384-well flat bottom, low volume microplate (Corning), and the fluorescence intensities were measured in a Synergy NEO2 plate reader (Biotek). Single turnover measurements were performed in the AutoSF 120 stopped-flow fluorimeter (Kintek) using 2x reconstituted holoenzyme with oPA (3 mM final) and 2x substrate, exciting at 550 nm and measuring Cy3 and Cy5 emission channels simultaneously.

The steady-state levels of FRET were calculated by the following equation: I_acceptor_/(I_donor_+I^−^_acceptor_). This value was calculated for each replicate and then averaged to determine the FRET signal for the respective condition. The FRET levels were then all normalized to that of the proteasome in ATP with no other additives. The kinetics of the conformational change were determined by fitting the gain of FRET observed in the Cy5 channel.

### Deubiquitination assays

Deubiquitination was tracked using FRET between Cy3-labeled ubiquitin and a Cy5-labeled substrate. Sulfo-Cy3 labeled ubiquitin was used to ubiquitinate Sulfo-Cy5 labeled substrates in the same ubiquitination conditions as described above. Single-turnover degradation reactions were then performed as described above, but instead of tracking anisotropy, Cy3 was excited, and Cy3 and Cy5 emission were monitored. The kinetics of deubiquitination were determined by fitting the loss of FRET observed in the Cy5 channel.

### Competition assays

The degradation competition experiments were performed similarly to the multiple-turnover degradation assays described above. The 2x reconstituted holoenzyme was mixed with a 2x solution containing both the labeled substrate and unlabeled competitor. The final concentrations used were 50 nM core, 400 nM base, 600 nM lid, 750 nM Rpn10, 500 nM FAM-titin-I27^V15P^-35, and varying competitor concentration. The extent of inhibition was then calculated by comparison of the initial rates with and without the competitor.

### Stability measurement

The equilibrium stability of titin mutants was measured by monitoring the tryptophan fluorescence of the substrate after equilibration at 25 °C with different concentrations of urea(¥2). The fluorescence (excitation = 280 nm, emission = 325 nm) was monitored in a plate reader at 25 °C. The curves were fit to the following system of equations:

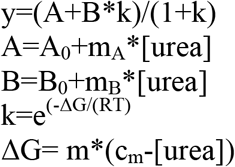

All variables were fit globally across the three substrates, except for the c_m_ values. The ΔG_0_ was then calculated using the equation ΔG_0_= m*c_m_.

### ATPase assays

The rate of ATP hydrolysis was measured using an NADH-coupled assay. Reconstituted proteasome (150 nM base, 400 nM core, 600 nM lid, 750 nM Rpn10, 1 mM ATP) was combined with ATPase mix (3 U ml^−1^ pyruvate kinase, 3 U ml^−1^ lactase dehydrogenase, 1 mM NADH, and 7.5 mM phosphoenolpyruvate), and the change of absorbance at 340 nm was monitored over time in a plate reader.

---

**Table S1.** Plasmids used for protein purification generated for this study

| Plasmid Name | Plasmid Description | Amino acid sequence after titin-I27* |
| --- | --- | --- |
| pAM81 | Rpn1, Rpn1, Rpn13 |  |
| pAM82 | 3xFLAG-Rpt1, Rpt2, 6xHis-Rpt3, Rpt4, Rpt5, Rpt6 |  |
| pAM83 | Nas2, Nas6, Hsm3, Rpn14, rare tRNAs |  |
| pAM87 | AzFRS.2.t1, UAG-tRNA |  |
| pAM88 | 3xFLAG-Rpt1I191TAG, Rpt2, 6xHis-Rpt3, Rpt4, Rpt5, Rpt6 |  |
| pAM89 | 3xFLAG-Rpt1, Rpt2, 6xHis-Rpt3, Rpt4, Rpt5Q49TAG, Rpt6 |  |
| pAM80 | Sem1, Hsp90 |  |
| pAM85 | Rpn5, MBP-HRV-Rpn6, Rpn8, Rpn9, Rpn11 |  |
| pAM86 | Rpn3, Rpn7, 6xHis-HRV-Rpn12 |  |
| pAM90 | Rpn5, MBP-HRV-Rpn6, Rpn8, Rpn9S111TAG, Rpn11 |  |
| pAM91 | GGG- titin-I27<sup>V15P</sup>-23-K-9-C-25-intein-CBD | GGAGPPPYSAANDENYALAAHGGKHTFNNE<br>NVSCRLGGAASIAVQAPAQHTFNNENVSY |
| pAM92 | GGG- titin-I27<sup>V15P</sup>-21-C-2-K-35-intein-CBD | GGAGPPPYSAANDENYALAAHCGGKHTFNN<br>ENVSARLGGAASIAVQAPAQHTFNNENVSY |
| pAM93 | GGG-titin-I27-23-K-9-C-25-intein-CBD | GGAGPPPYSAANDENYALAAHGGKHTFNNE<br>NVSCRLGGAASIAVQAPAQHTFNNENVSY |
| pAM94 | GGG- titin-I27<sup>V13P/V15P</sup>-23-K-9-C-25-intein-CBD | GGAGPPPYSAANDENYALAAHGGKHTFNNE<br>NVSCRLGGAASIAVQAPAQHTFNNENVSY |
| pAM95 | GGG- titin-I27<sup>V15P</sup>-43-K-8-C-25-intein-CBD | GGAGPPPYSAANDENYALAAHGGGAHTFNNE<br>NVSAHTFNNENVSKHTFNNENVCRLGGAASI<br>AVQAPAQHTFNNENVSY |
| pAM96 | GGG- titin-I27<sup>V15P</sup>-33-K-9-K-8-C-25 | GGAGPPPYSAANDENYALAAHGGGAHTFNNE<br>NVSKHTFNNENVSKHTFNNENVCRLGGAASI<br>AVQAPAQHTFNNENVSY |
| pAM97 | GGG- titin-I27<sup>V15P</sup>-23-K-9-K-9-K-8-C-25 | GGAGPPPYSAANDENYALAAHGGKHTFNNE<br>NVSKHTFNNENVSKHTFNNENVCRLGGAASI<br>AVQAPAQHTFNNENVSY |
| pAM98 | titin-I27<sup>V15P</sup>-23-K-9-C-15 | GGAGPPPYSAANDENYALAAHGGKHTFNNE<br>NVSCRLGGAASIAVQAPAY |
| pAM99 | titin-I27<sup>V15P</sup>-23-K-9-C-1 | GGAGPPPYSAANDENYALAAHGGKHTFNNE<br>NVSCY |
| pAM100 | titin-I27<sup>V15P</sup>-21-C-2-K-1 | GGAGPPPYSAANDENYALAAHCGGKY |
| pAM101 | titin-I27<sup>V15P</sup>-23-K-9-C-15 serine rich | GGAGPPPYSAANDENYALAAHGGKSSSSSSA<br>SSCSSGSSSSSSASSSSY |
| pAM102 | 6xHis-Sumo-Ub-Ub-Ub-Ub |  |
| pAM103 | MC-Ubiquitin |  |

\*The amino acid sequence of the lysine-less version of titin-I27 used is as follows:
LIEVERPLYGVEVFGVGETAHFEIELSEPDVHGQWRLRGQPLAASPDCEIIEDGRRHILILHNCQLGMTGEVSF QAANTRSAANLRVREL

**Table S2.** Substrates used for each assay

| Plasmid Origin | Description | N-terminal modification | Tail modification | Figure |
| --- | --- | --- | --- | --- |
| pAM91 | titin-I27<sup>V15P</sup>-35 | - | - | 1D, 2E-F, 4C-D, S4B, S6, Table S5, Table S7 |
| pAM91 | titin-I27<sup>V15P</sup>-35 | 5-FAM | - | 2B-C, 3A-B, 4D, S1-3, S4D, S7C, S10, Table S3 |
| pAM91 | titin-I27<sup>V15P</sup>-35 | - | 5-FAM | 4A, S9 |
| pAM91 | titin-I27<sup>V15P</sup>-35 | 5-FAM | Cy5 | Table S3 |
| pAM91 | titin-I27<sup>V15P</sup>-35 | - | Cy5 | 2D, 4B, S4A, Table S4 |
| pAM92 | titin-I27<sup>V15P</sup>-35 (DUB) | 5-FAM | Cy5 | 2G, S4C, S5, Table S6 |
| pAM93 | titin-I27-35 | - | - | S6, Table S7 |
| pAM93 | titin-I27-35 | 5-FAM | - | 3A, 3B, Table S3 |
| pAM93 | titin-I27-35 | 5-FAM | Cy5 | Table S3 |
| pAM94 | titin-I27<sup>V13P/V15P</sup>-35 | - | - | S6, Table S7 |
| pAM94 | titin-I27<sup>V13P/V15P</sup>-35 | 5-FAM | - | 3A, 3B, Table S3 |
| pAM95 | titin-I27<sup>V15P</sup>-1K | 5-FAM | - | 3C, S2, S7, Table S3 |
| pAM96 | titin-I27<sup>V15P</sup>-2K | 5-FAM | - | 3C, S2, S7, Table S3 |
| pAM97 | titin-I27<sup>V15P</sup>-3K | 5-FAM | - | 3C, S2, S7, Table S3 |
| pAM98 | titin-I27<sup>V15P</sup>-25 | - | Cy5 | 4B, Table S4 |
| pAM98 | titin-I27<sup>V15P</sup>-25 | - | 5-FAM | Table S3, S8 |
| pAM99 | titin-I27<sup>V15P</sup>-11 | - | Cy5 | 4B, Table S4 |
| pAM99 | titin-I27<sup>V15P</sup>-11 | - | - | 4C, 4D |
| pAM99 | titin-I27<sup>V15P</sup>-11 | - | 5-FAM | 4A, S9, S11 |
| pAM100 | titin-I27<sup>V15P</sup>-1 | - | Cy5 | 4B, Table S4 |
| pAM100 | titin-I27<sup>V15P</sup>-1 | - | - | 4C, 4D |
| pAM100 | titin-I27<sup>V15P</sup>-1 | - | 5-FAM | 4A, S2, S9, S11 |
| pAM101 | titin-I27<sup>V15P</sup>-serine rich | - | Cy5 | 4B, Table S4 |
| pAM101 | titin-I27<sup>V15P</sup>-serine rich | - | - | 4C, 4D |
| pAM101 | titin-I27<sup>V15P</sup>-serine rich | - | 5-FAM | 4A, S8-9, S11 |

**Table S3.** Degradation fits

| Substrate Description | $A_1/A_2$ | $\tau_1+t_0$ (s) | $\tau_2+t_0$ (s) | $t_0$ (s) |
| --- | --- | --- | --- | --- |
| FAM- titin-I27-35 | $2.3 \pm 0.3$ | $47.2 \pm 2.1$ | $499 \pm 98$ | 11.0 |
| FAM-titin-I27-35-Cy5 | $4.2 \pm 0.3$ | $45.0 \pm 1.8$ | $223 \pm 18$ | 11.5 |
| FAM-titin-I27<sup>V15P</sup>-35 | $2.1 \pm 0.2$ | $18.1 \pm 0.4$ | $89 \pm 10$ | 7.4 |
| FAM-titin-I27<sup>V15P</sup>-35-Cy5 | $3.0 \pm 0.6$ | $20.0 \pm 0.7$ | $139 \pm 44$ | 7.5 |
| FAM-titin-I27<sup>V13P/V15P</sup>-35 | $1.7 \pm 0.1$ | $14.7 \pm 0.3$ | $77 \pm 5$ | 6.0 |
| titin-I27<sup>V15P</sup>-25-FAM | $5.4 \pm 0.5$ | $21.1 \pm 0.5$ | $157 \pm 30$ | 7.5 |
| FAM-titin-I27<sup>V15P</sup>-1k | $2.6 \pm 0.5$ | $22.0 \pm 1.8$ | $111 \pm 22$ | 10.0 |
| FAM-titin-I27<sup>V15P</sup>-2k | $3.1 \pm 0.7$ | $21.0 \pm 0.8$ | $108 \pm 14$ | 10.0 |
| FAM-titin-I27<sup>V15P</sup>-3k | $3.2 \pm 0.8$ | $21.4 \pm 0.2$ | $126 \pm 12$ | 10.0 |

**Table S4.**
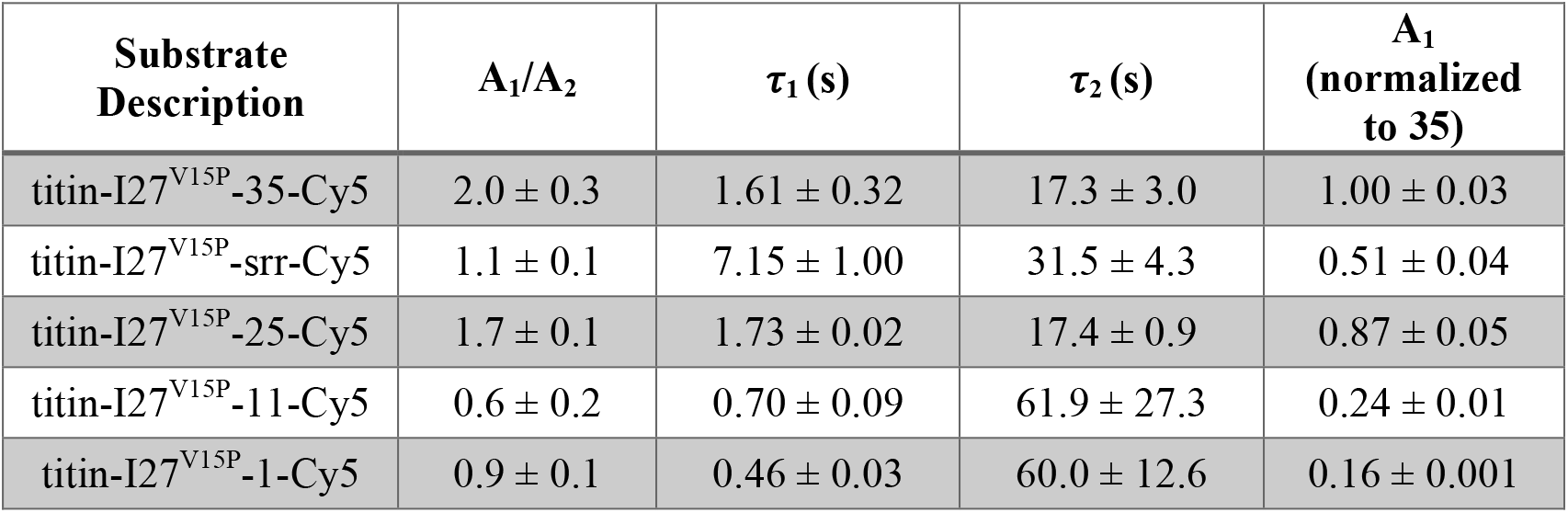
Tail insertion fits

| Substrate Description | $A_1/A_2$ | $\tau_1$ (s) | $\tau_2$ (s) | $A_1$ (normalized to 35) |
| --- | --- | --- | --- | --- |
| titin-I27<sup>V15P</sup>-35-Cy5 | $2.0 \pm 0.3$ | $1.61 \pm 0.32$ | $17.3 \pm 3.0$ | $1.00 \pm 0.03$ |
| titin-I27<sup>V15P</sup>-srr-Cy5 | $1.1 \pm 0.1$ | $7.15 \pm 1.00$ | $31.5 \pm 4.3$ | $0.51 \pm 0.04$ |
| titin-I27<sup>V15P</sup>-25-Cy5 | $1.7 \pm 0.1$ | $1.73 \pm 0.02$ | $17.4 \pm 0.9$ | $0.87 \pm 0.05$ |
| titin-I27<sup>V15P</sup>-11-Cy5 | $0.6 \pm 0.2$ | $0.70 \pm 0.09$ | $61.9 \pm 27.3$ | $0.24 \pm 0.01$ |
| titin-I27<sup>V15P</sup>-1-Cy5 | $0.9 \pm 0.1$ | $0.46 \pm 0.03$ | $60.0 \pm 12.6$ | $0.16 \pm 0.001$ |

**Table S5.** Conformational change fit

| Substrate Description | $A_1/A_2$ | $\tau_1+t_0$ (s) | $\tau_2+t_0$ (s) | $t_0$ (s) |
| --- | --- | --- | --- | --- |
| titin-I27<sup>V15P</sup>-35 | $2.3 \pm 0.2$ | $2.22 \pm 0.10$ | $24.9 \pm 1.8$ | 0.4 |

**Table S6.**
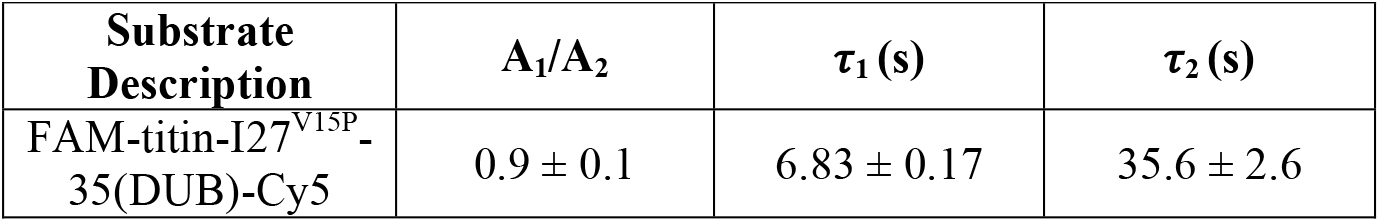
Deubiquitination fit

**Table S7.** ATPase rates

| Sample | ATPase rate (ATP s<sup>-1</sup> enzyme<sup>-1</sup>) |
| --- | --- |
| 26S alone | $0.73 \pm 0.02$ |
| 26S + ub'd titin-I27<sup>V13P/V15P</sup>-35 | $1.12 \pm 0.11$ |
| 26S + ub'd titin-I27<sup>V15P</sup>-35 | $1.35 \pm 0.12$ |
| 26S + ub'd titin-I27-35 | $1.47 \pm 0.11$ |

**Fig. S1.**
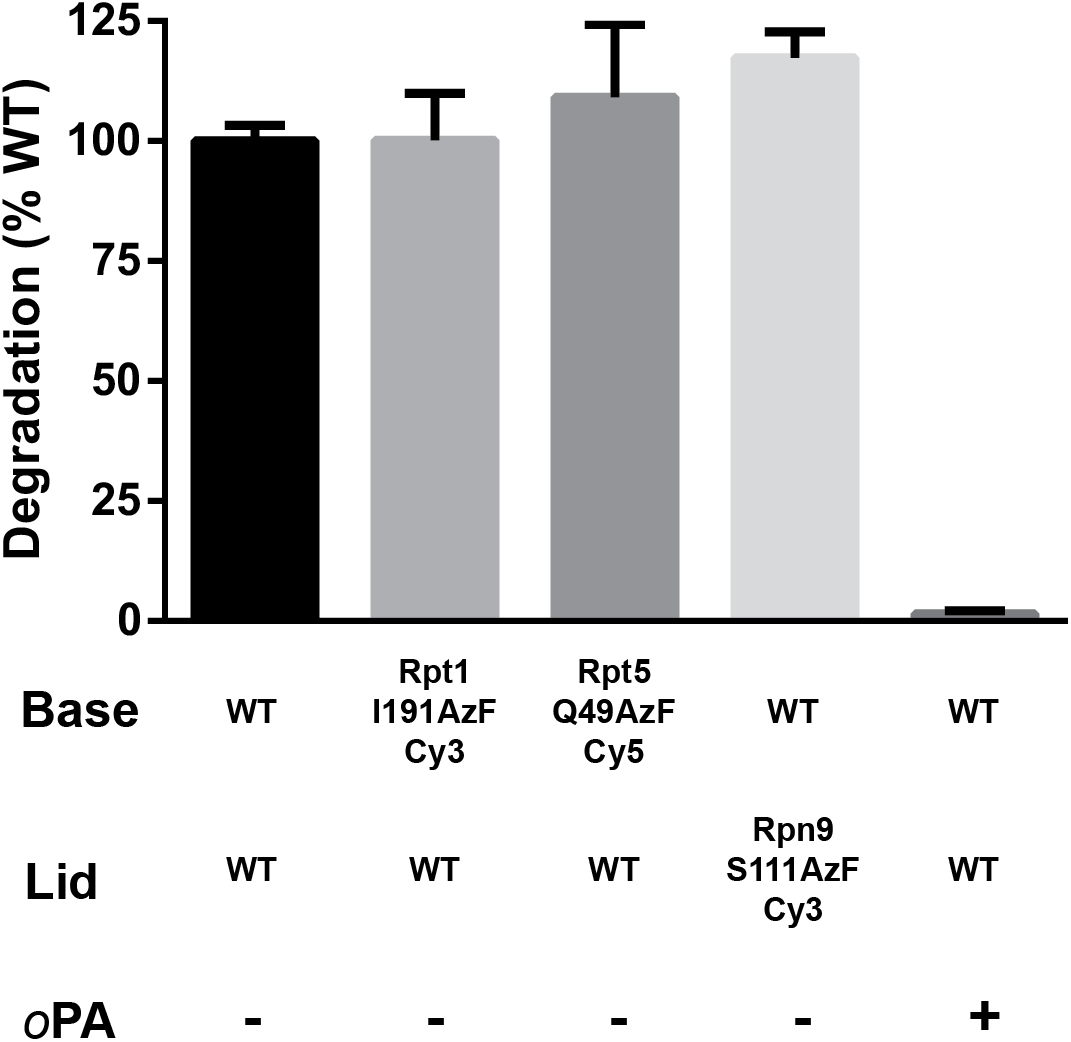
Multiple-turnover degradations with labeled components. Initial rates for multiple-turnover degradations of FAM-titin-I27^V15P^-35 by 26S proteasome, tracked by anisotropy of 5-FAM covalently attached to the N-terminus of the substrate. Proteasomes were reconstituted using the indicated lid and base components, limiting 20S core, and Rpn10. Degradation rates are normalized to that of proteasomes reconstituted with all wild-type components. 1,10-phenanthroline (oPA) inhibits Rpn11 ubiquitin cleavage and thus prevents degradation. Data shown are means and s.d. (N=3).

**Fig. S2.**
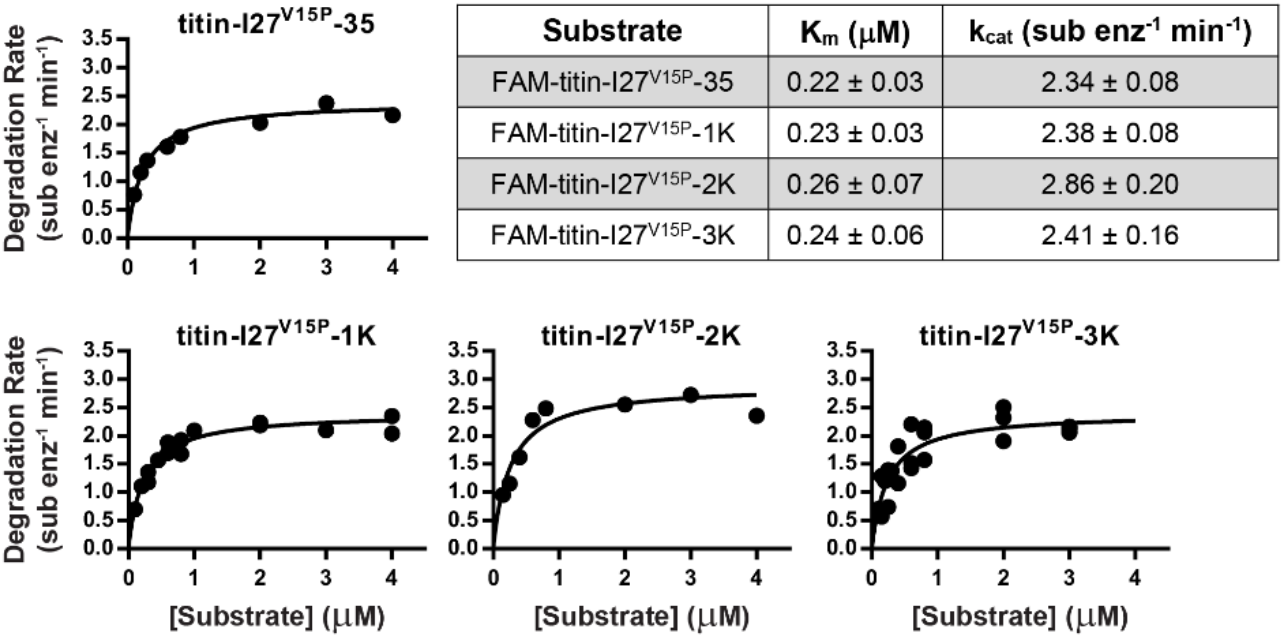
Michaelis Menten analysis of substrate variants. The initial rates for multiple-turnover degradation of the indicated substrates at varying concentrations were determined by tracking anisotropy of 5-FAM attached to the N-terminus of the titin-127 domain. The curves were fit to the Michaelis-Menten equation, and K_M_ and k_cat_ are reported with the standard error of the fits.

**Fig. S3.**
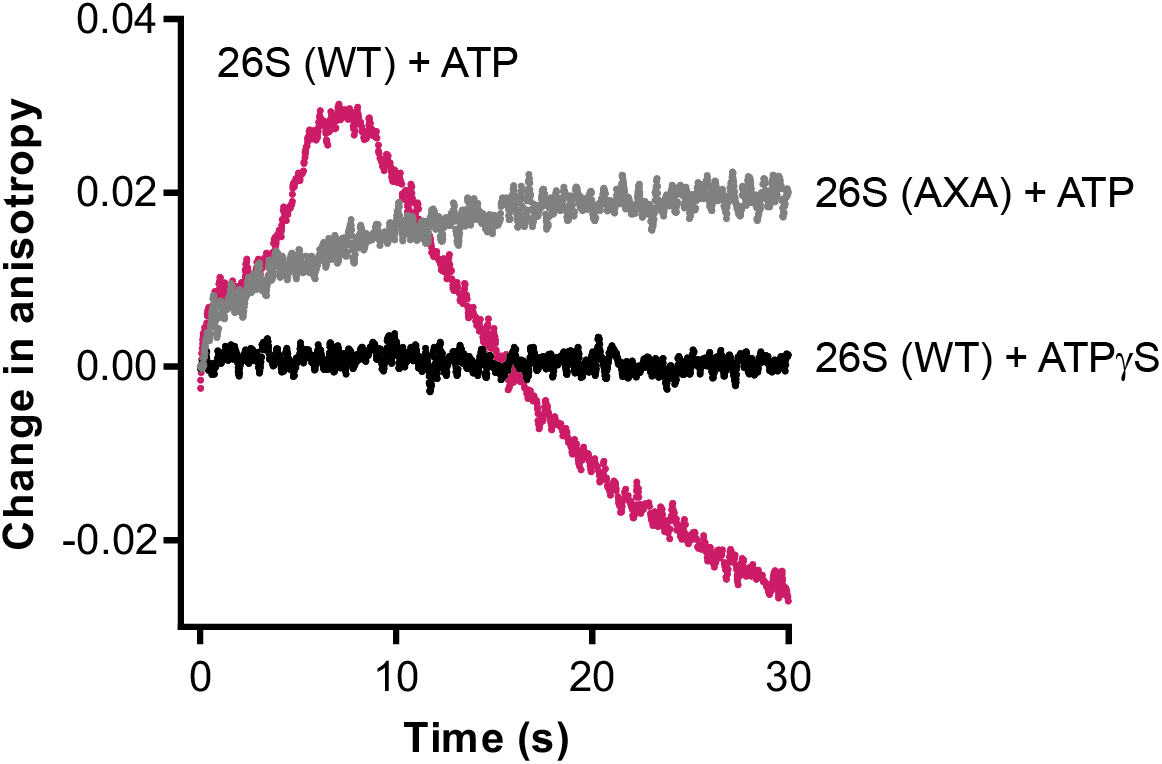
Comparison of AXA vs. WT 26S. The anisotropy change of FAM-titin-I27^V15P^-35 after the addition of reconstituted 26S proteasome in the presence of either ATP or ATPgS. The 26S proteasome was reconstituted with either wild-type (WT) or catalytically-dead Rpn11^AXA^ (AXA) lid.

**Fig. S4.**
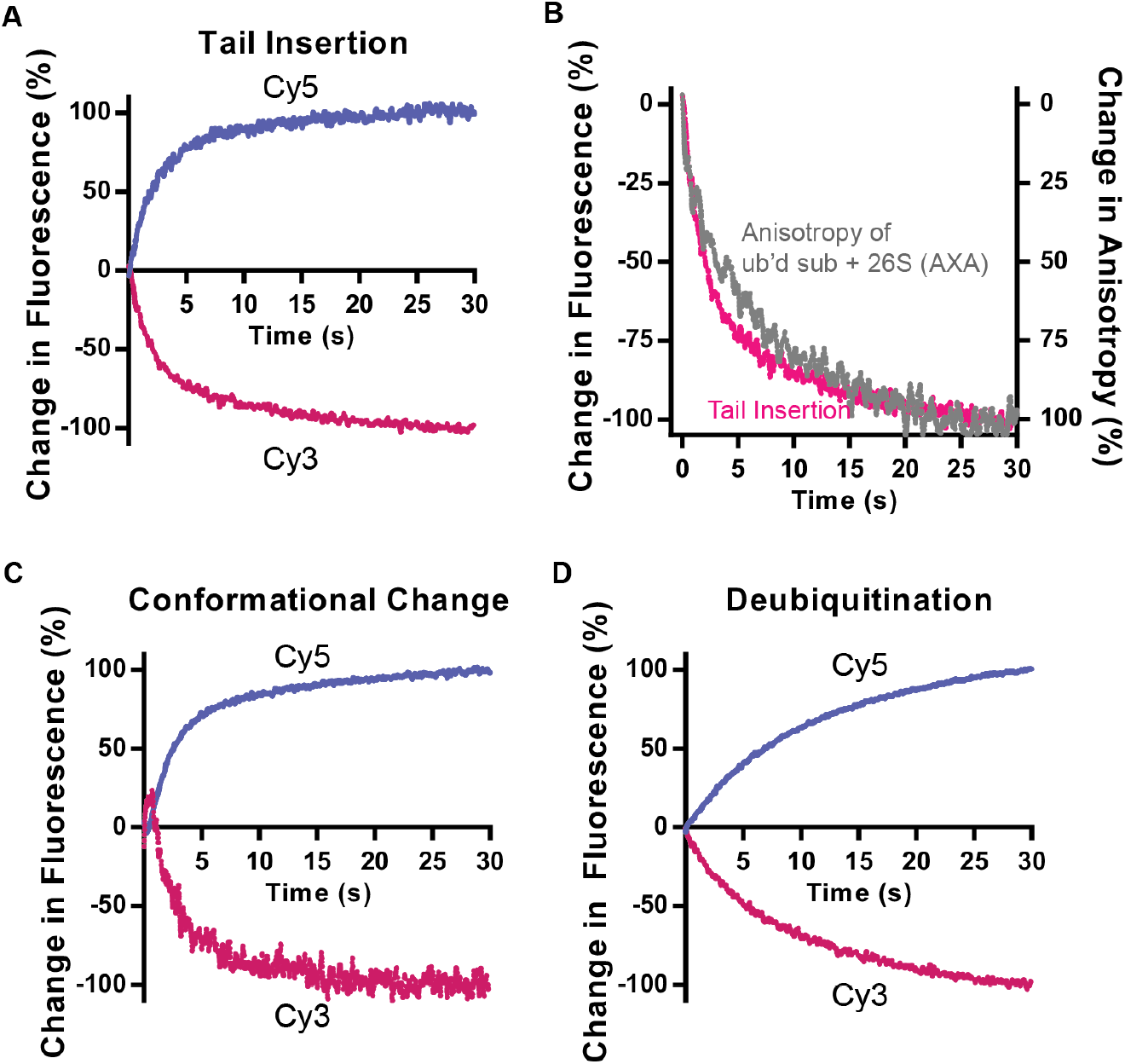
Reciprocal relationship of Cy3 and Cy5 fluorescence during FRET experiments. Representative traces of each fluorescence channel from the FRET experiments in Fig. 2. Each trace was normalized to both the initial fluorescence (0%) and the fluorescence after 30 s (100%). **(A)** Ubiquitinated titin-I27^v15P^-35-Cy5 was mixed with oPA-treated, Rpt1-I191AzF-Cy3-containing proteasome in the presence of ATP, as in Fig. 2D. **(B)** Overlay of the Cy3 channel from **A** and the anisotropy of FAM-titin-I27^v15P^-35 after mixing with 26S proteasome containing catalytically dead Rpn11^AXA^ (AXA), as in Fig. S3. Like the fluorescence channel, the anisotropy trace was normalized to both the anisotropy at t=0 (0%) and the anisotropy after 30 s (100%). **(C)** oPA-treated proteasome containing Rpn9S111AzF-Cy3 and Rpt5Q49AzF-Cy5, as in Fig. 2E, were mixed with ubiquitinated titin-I27^v15P^-35 and ATP. **(D)** Titin-I27^v15P^-35-Cy5 ubiquitinated with Cy3-ubiquitin as in Fig. 2G, was mixed with proteasome and ATP.

**Fig. S5.**
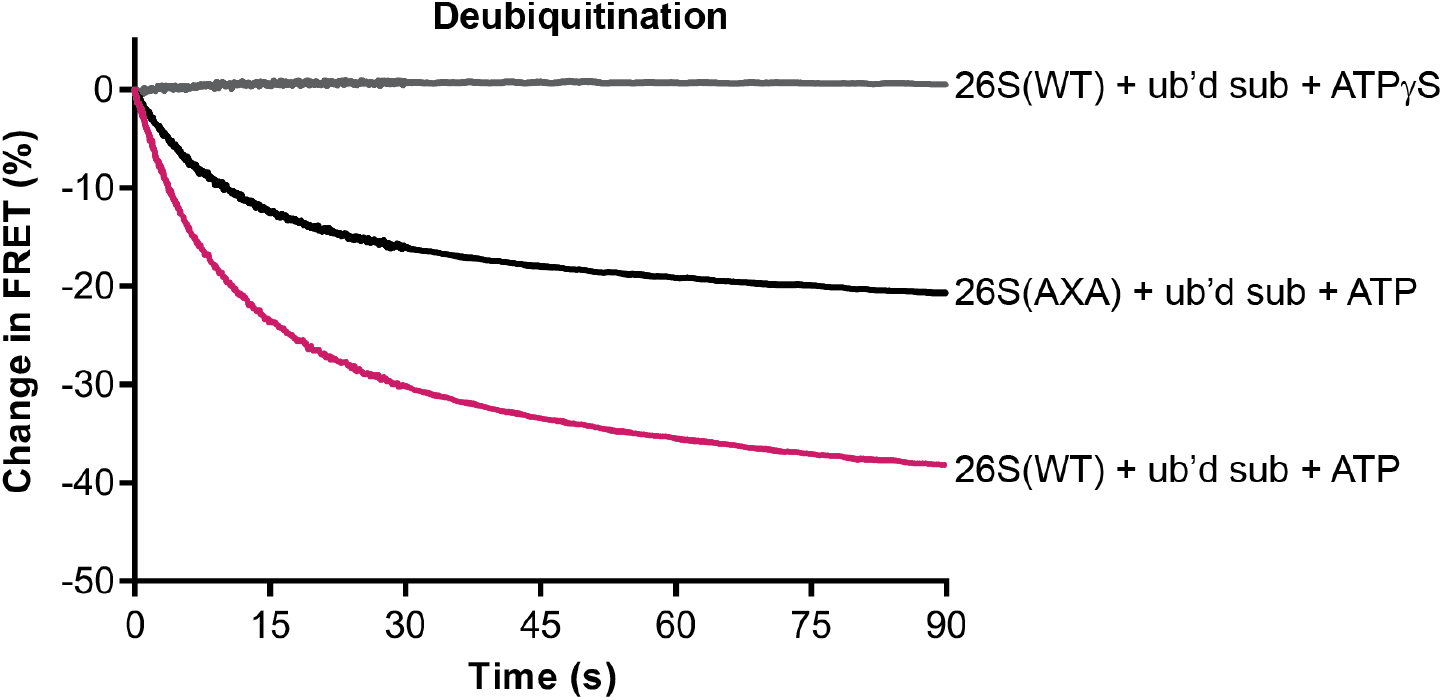
FRET between ubiquitin and substrate. Titin-I27^v15P^-35-Cy5 substrate modified with Cy3-labeled ubiquitin was mixed with wild-type 26S proteasome in the presence of ATP or ATPγS, or with proteasome reconstituted with Rpn11^AXA^ lid, and monitored by Cy5 fluorescence.

**Fig. S6.**
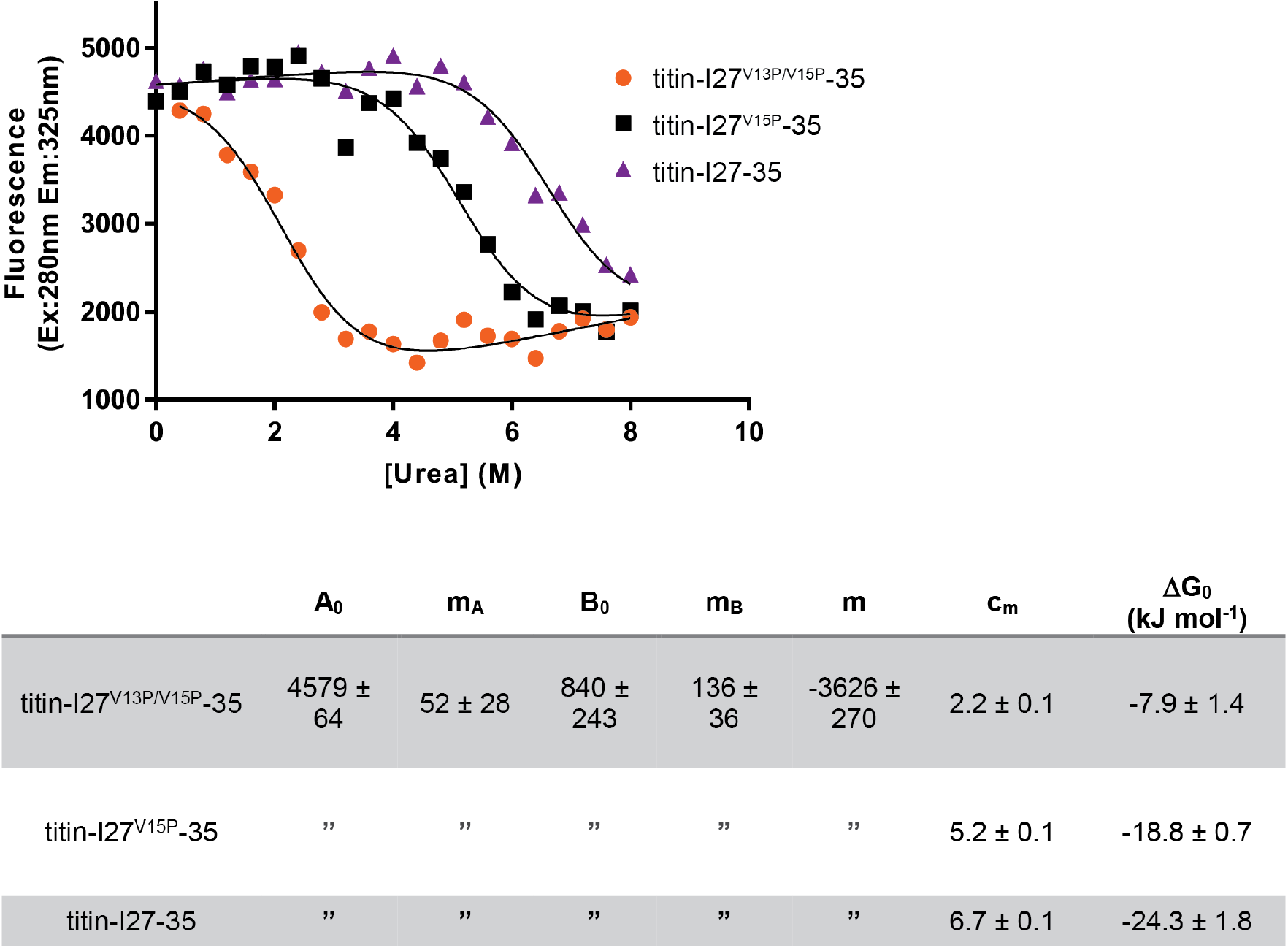
Determining the thermodynamic stability of titin variants. Denaturant-induced equilibrium unfolding of substrates with different titin stabilities. The folded state of titin-127 was monitored by tracking intrinsic tryptophan fluorescence. The curves were fit to the following system of equations: y=(A+B*k)/(1+k), A=A_0_+m_A_*[urea], B=B_0_+m_B_*[urea], k=e^(-ΔG/(RT)^, ΔG= m*(c_m_-[urea]). All variables were fit globally across the three substrates, except for the c_m_ values. The ΔGo was then calculated using the equation ΔGo= m*c_m_.

**Fig. S7.**
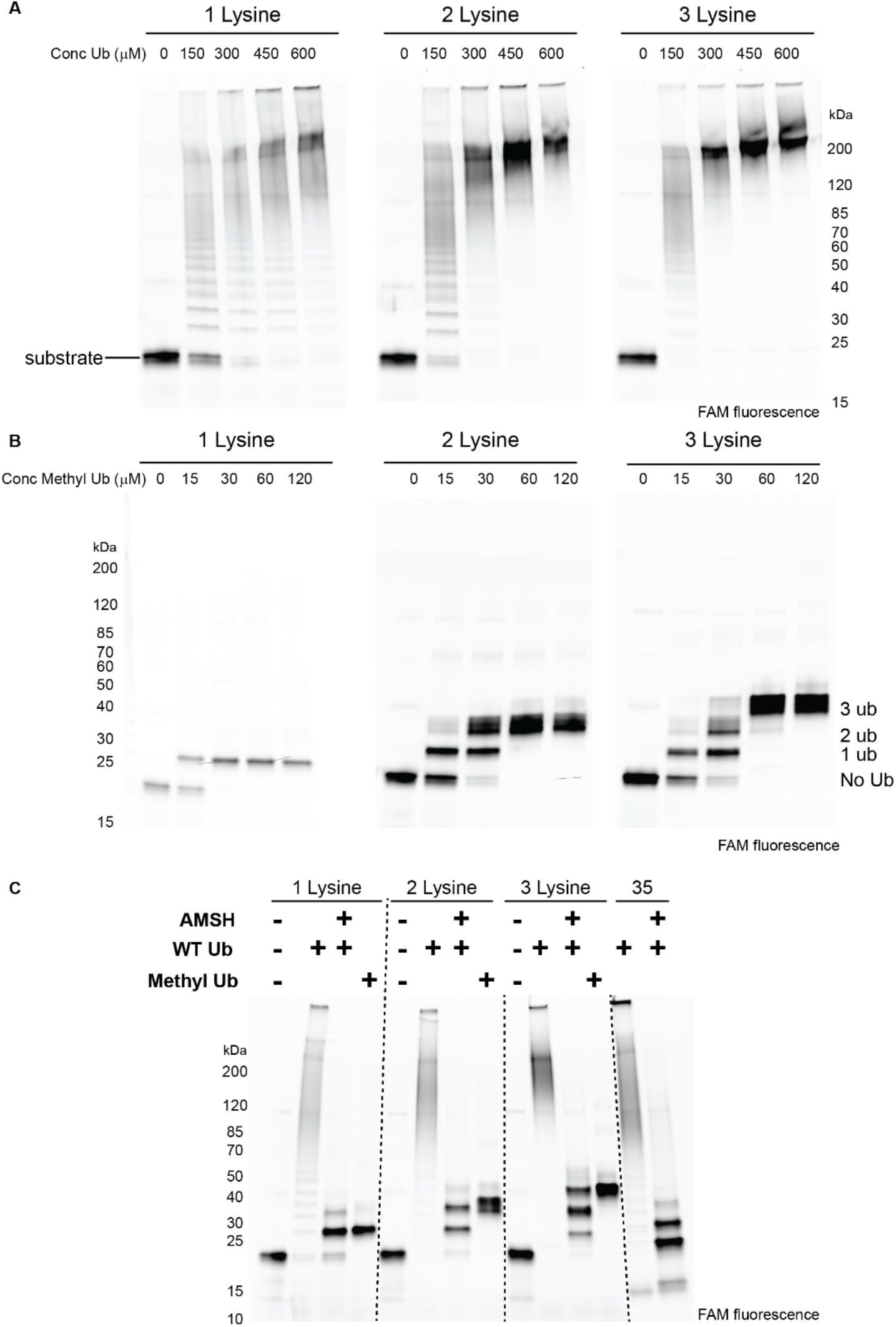
Ubiquitination state of multiple lysine substrates. Substrates with 1, 2, or 3 lysines were ubiquitinated with varying concentrations of ubiquitin, analyzed by SDS-PAGE and visualized by 5-FAM fluorescence. Wild-type ubiquitin is used in **(A)**,methyl ubiquitin, which is unable to form chains, is used in **(B)**. In **(C),** the substrates were modified with either 400 μM wild-type ubiquitin or 120 μM methyl ubiquitin. The sample with wild-type ubiquitin was treated with the K63-specific deubiquitinase AMSH. Also in **(C),** the FAM-titin-I27^V15P^-35 was ubiquitinated with 400 μM wild-type ubiquitin and treated with AMSH.

**Fig. S8.**
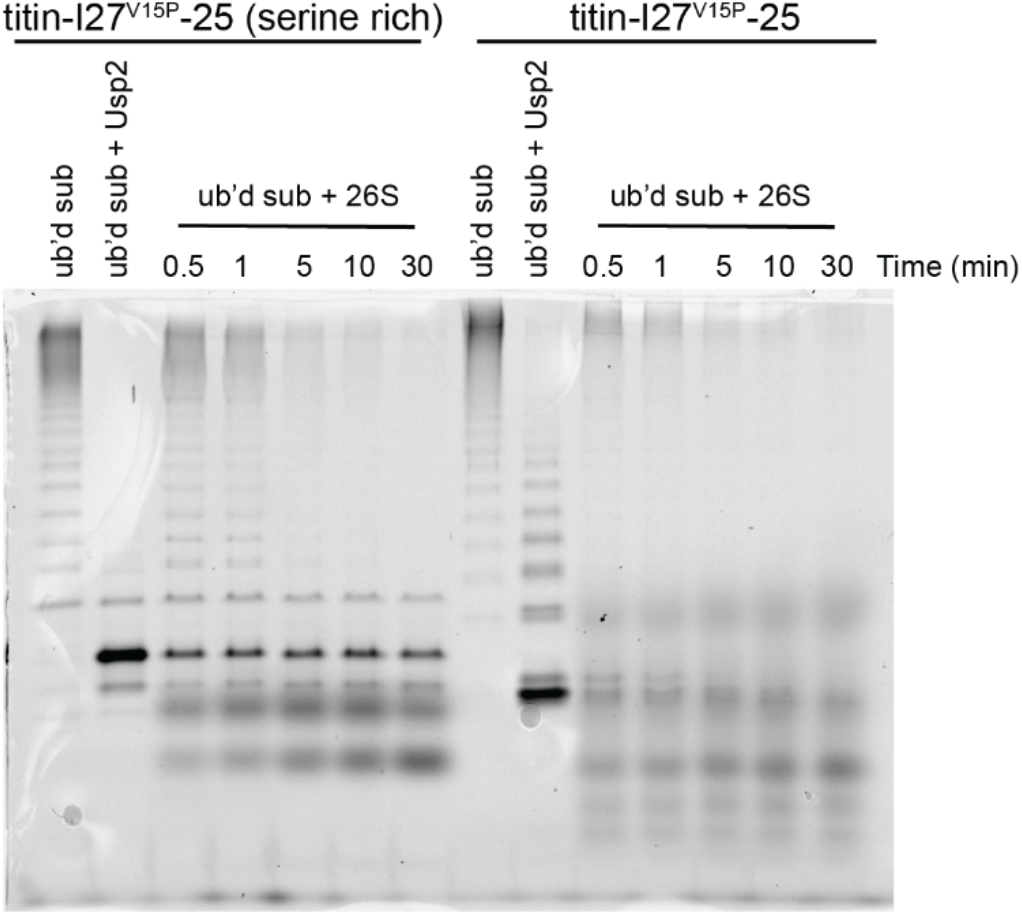
Comparison of substrates with 25 amino acid tails. Single-turnover degradations of substrates with 25 amino acid tails, labeled with 5-FAM on the C-terminal tail, analyzed by SDS-PAGE, and visualized by 5-FAM fluorescence. Also shown is a control with the ubiquitinated substrate after addition of the non-specific deubiquitinase Usp2.

**Fig. S9.**
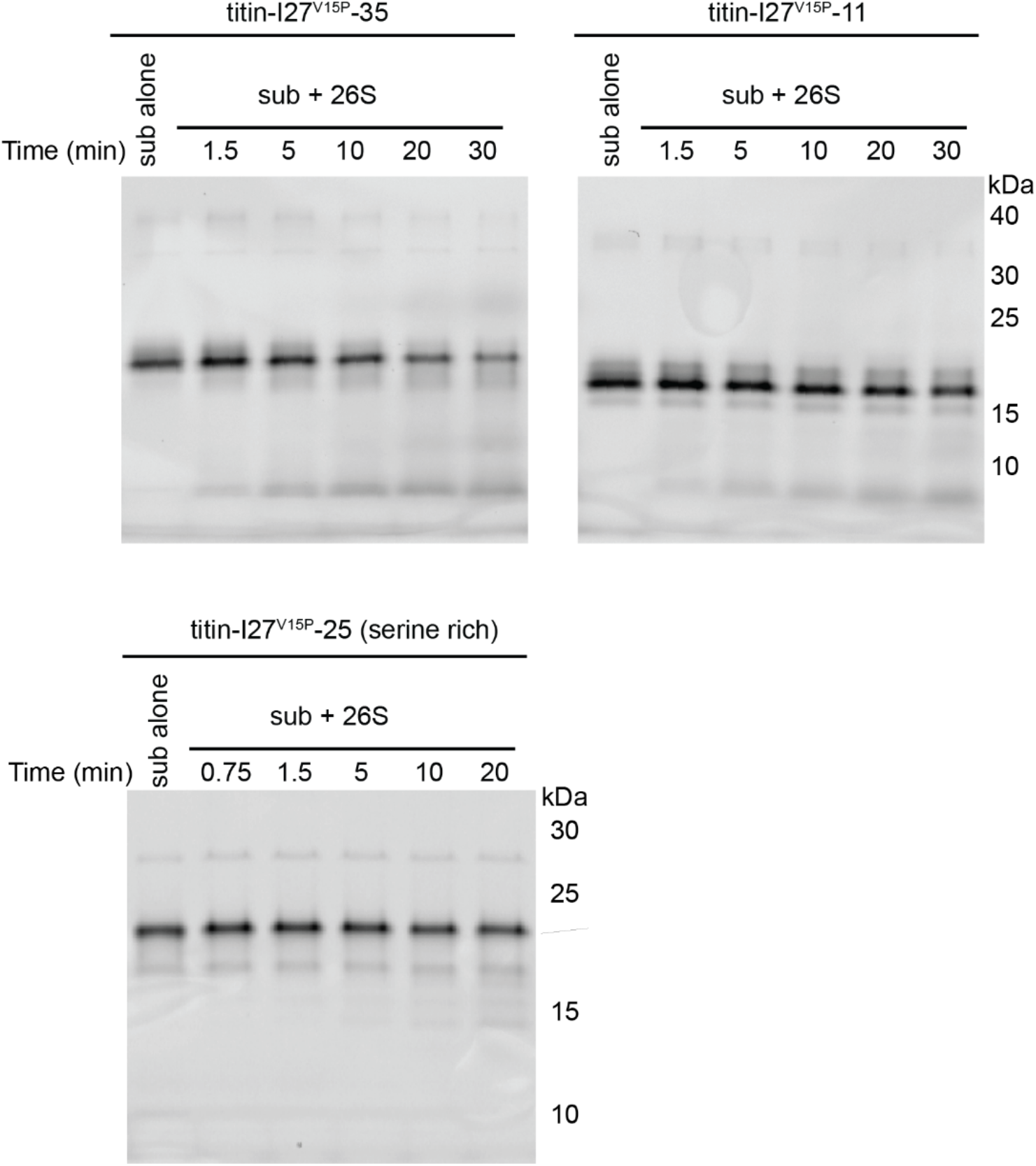
Non-ubiquitinated substrate degradation. Single-turnover degradations of non-ubiquitinated substrates, labeled with 5-FAM on the C-terminal tail, using the same reconstituted proteasome conditions as those used in Fig. 4D. The degradation observed may be a result of unbound 20S core particle directly proteolyzing substrates and could account for the small accumulation of peptide products observed for titin-I27^V15^-11 in Fig. 4A.

**Fig. S10.**
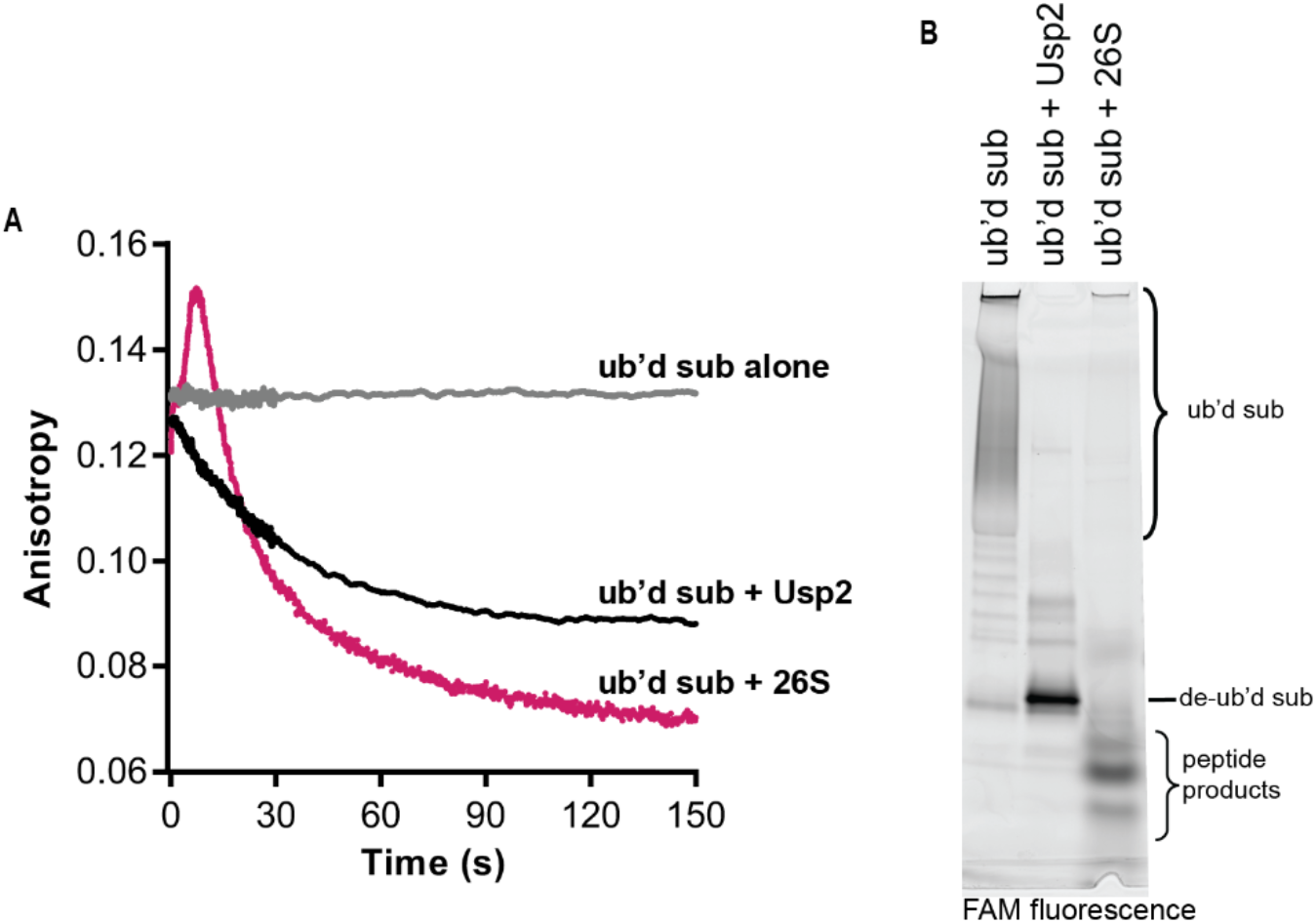
Anisotropy can be used to track deubiquitination. **(A)** Anisotropy change of FAM-titin-I27^v15P^-35 after addition of reconstituted 26S proteasome with ATP or addition of the non-specific deubiquitinase Usp2. The anisotropy change is greater after addition of proteasome due to proteolytic cleavage of the substrate into small peptides. **(B)** SDS-PAGE analysis of ubiquitinated FAM-titin-I27^v15P^-35 alone, after addition of Usp2 or 26S proteasome.

**Fig. S11.**
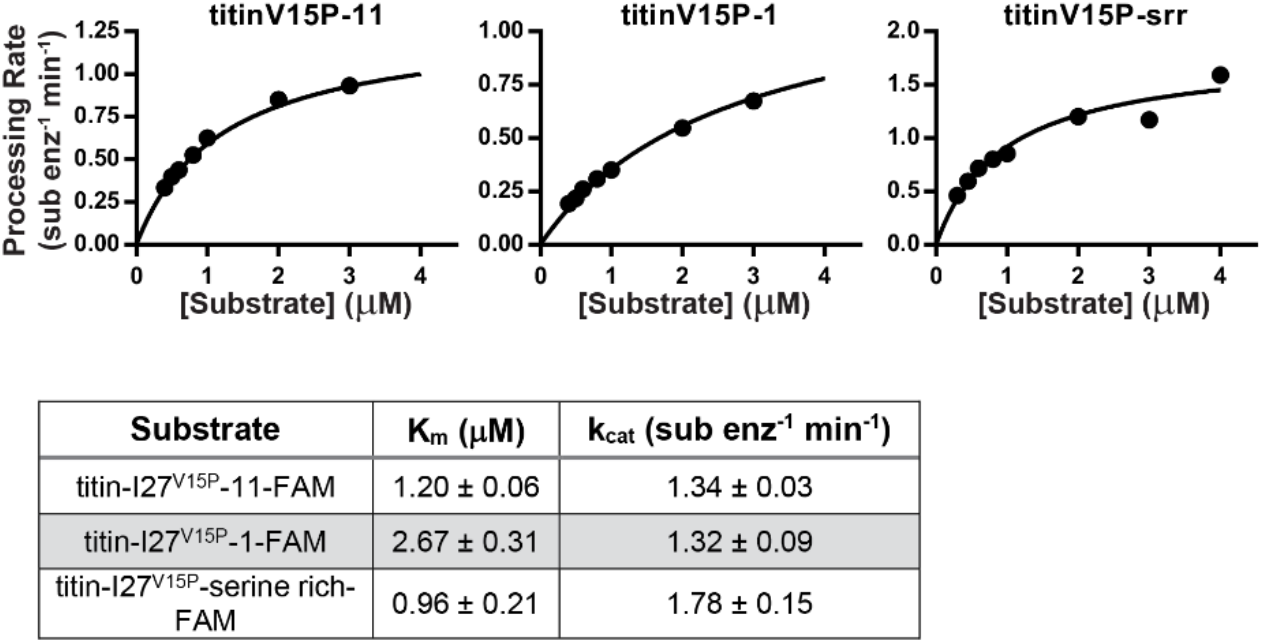
Michaelis-Menten analyses of substrate variants. The initial rates of multiple-turnover processing of the indicated substrates at varying concentrations were determined by tracking the anisotropy of 5-FAM attached to the C-terminal unstructured region. The curves were fit to the Michaelis-Menten equation, and K_M_ and k_cat_ are reported with the standard error of the fits.

